# Widespread redundancy in -omics profiles of cancer mutation states

**DOI:** 10.1101/2021.10.27.466140

**Authors:** Jake Crawford, Brock C. Christensen, Maria Chikina, Casey S. Greene

**Affiliations:** Genomics and Computational Biology Graduate Group, Perelman School of Medicine, University of Pennsylvania, Philadelphia, PA, USA; Funded by National Institutes of Health’s National Cancer Institute (R01 CA237170); National Institutes of Health’s National Human Genome Research Institute(R01 CA216265, R01 CA253976); Department of Epidemiology, Geisel School of Medicine, Dartmouth College, Lebanon, NH, USA; Department of Molecular and Systems Biology, Geisel School of Medicine, Dartmouth College, Lebanon, NH, USA; Funded by National Institutes of Health’s National Cancer Institute (R01 CA216265, R01 CA253976); Department of Computational and Systems Biology, School of Medicine, University of Pittsburgh, Pittsburgh, PA, USA; Department of Biochemistry and Molecular Genetics, University of Colorado School of Medicine, Aurora, CO, USA; Center for Health AI, University of Colorado School of Medicine, Aurora, CO, USA; Funded by National Institutes of Health’s National Cancer Institute (R01 CA237170); National Institutes of Health’s National Human Genome Research Institute(R01 HG010067)

## Abstract

In studies of cellular function in cancer, researchers are increasingly able to choose from many -omics assays as functional readouts. Choosing the correct readout for a given study can be difficult, and which layer of cellular function is most suitable to capture the relevant signal may be unclear. In this study, we consider prediction of cancer mutation status (presence or absence) from functional -omics data as a representative problem. Since functional signatures of cancer mutation have been identified across many data types, this problem presents an opportunity to quantify and compare the ability of different -omics readouts to capture signals of dysregulation in cancer. The TCGA Pan-Cancer Atlas contains genetic alteration data including somatic mutations and copy number variants (CNVs), as well as several -omics data types. From TCGA, we focus on RNA sequencing, DNA methylation arrays, reverse phase protein arrays (RPPA), microRNA, and somatic mutational signatures as -omics readouts.

Across a collection of genes recurrently mutated in cancer, RNA sequencing tends to be the most effective predictor of mutation state. Surprisingly, we found that for many of the genes we considered, one or more other data types are approximately equally effective predictors. Performance was more variable between mutations than it was between data types for the same mutation, and there was often little difference between the top data types. We also found that combining data types into a single multi-omics model provided little or no improvement in predictive ability over the best individual data type. Based on our results, for the design of studies focused on the functional outcomes of cancer mutations, there are often multiple -omics types that can serve as effective readouts, although gene expression seems to be a reasonable default option.

## Introduction

Although cancer can be initiated and driven by many different genetic alterations, these tend to converge on a limited number of pathways or signaling processes [1]. As driver mutation status alone confers limited prognostic information, a comprehensive understanding of how diverse genetic alterations perturb central pathways is vital to precision medicine and biomarker identification efforts [2,3]. While many methods exist to distinguish driver mutations from passenger mutations based on genomic sequence characteristics [4,5,6], until recently it has been a challenge to connect driver mutations to downstream changes in gene expression and cellular function within individual tumor samples.

The Cancer Genome Atlas (TCGA) Pan-Cancer Atlas provides uniformly processed, multi-platform - omics measurements across tens of thousands of samples from 33 cancer types [7]. Enabled by this publicly available data, a growing body of work on linking the presence of driving genetic alterations in cancer to downstream gene expression changes has emerged. Recent studies have considered Ras pathway alteration status in colorectal cancer [8], alteration status across many cancer types in Ras genes [9,10], *TP53* [11], and *PIK3CA* [12], and alteration status across cancer types in frequently mutated genes [13]. More broadly, other groups have drawn on similar ideas to distinguish between the functional effects of different alterations in the same driver gene [14], to link alterations with similar gene expression signatures within cancer types [15], and to identify trans-acting expression quantitative trait loci (trans-eQTLs) in germline genetic studies [16].

These studies share a common thread: they each combine genomic (point mutation and copy number variation) data with transcriptomic (RNA sequencing) data within samples to interrogate the functional effects of genetic variation. RNA sequencing is ubiquitous and cheap, and its experimental and computational methods are relatively mature, making it a vital tool for generating insight into cancer pathology [17]. Some driver mutations, however, are known to act indirectly on gene expression through varying mechanisms. For example, oncogenic *IDH1* and *IDH2* mutations in glioma have been shown to interfere with histone demethylation, which results in increased DNA methylation and blocked cell differentiation [18,19,20,21]. Other genes implicated in aberrant DNA methylation in cancer include the TET family of genes [22] and *SETD2* [23]. Certain driver mutations, such as those in DNA damage repair genes, may lead to detectable patterns of somatic mutation [24]. Additionally, correlation between gene expression and protein abundance in cancer cell lines is limited, and proteomics data could correspond more directly to certain cancer phenotypes and pathway perturbations [25]. In these contexts and others, integrating different data modalities or combining multiple data modalities could be more effective than relying solely on gene expression as a functional signature.

Here, we compare -omics data types profiled in the TCGA Pan-Cancer Atlas to evaluate use as a multivariate functional readout of genetic alterations in cancer. We focus on gene expression (RNA sequencing data), DNA methylation (27K and 450K probe chips), reverse phase protein array (RPPA), microRNA expression, and mutational signatures data [26] as possible readouts. Prior studies have identified univariate correlations of CpG site methylation [27,28] and correlations of RPPA protein profiles [29] with the presence or absence of certain driver mutations. Other relevant past work includes linking point mutations and copy number variants (CNVs) with changes in methylation and expression at individual genes [30,31] and identifying functional modules that are perturbed by somatic mutations [32,33]. However, direct comparison among different data types for this application is lacking, particularly in the multivariate case where we consider changes to -omics-derived gene signatures rather than individual genes in isolation.

We select a collection of potential cancer drivers with varying functions and roles in cancer development. We use mutation status in these genes as labels to train classifiers, using each of the data types listed as training data, in a pan-cancer setting; we follow similar methods to the elastic net logistic regression approach described in Way et al. 2018 [9] and Way et al. 2020 [13]. We show that there is considerable predictive signal for many genes relative to a cancer-type corrected baseline and that gene expression tends to provide good predictions of mutation state across most genes.

Surprisingly, we find that for a variety of genes, multiple data types are approximately equally effective predictors. We observe similar results for pan-cancer survival prediction across the same data types with little separation between the top-performing data types. In addition, we observe that combining data types into a single multi-omics model for mutation prediction provides little, if any, performance benefit over the most performant model using a single data type. Our results will help to inform the design of future functional genomics studies in cancer, suggesting that for many strong drivers with clear functional signatures, different -omics measurements can provide similar information content.

## Methods

### Mutation data download and preprocessing

To generate binary mutated/non-mutated gene labels for our machine learning model, we used mutation calls for TCGA samples from MC3 [34] and copy number threshold calls from GISTIC2.0 [35]. MC3 mutation calls were downloaded from the Genomic Data Commons (GDC) of the National Cancer Institute, at https://gdc.cancer.gov/about-data/publications/pancanatlas. Copy number threshold calls are from an older version of the GDC data, and are available here: https://figshare.com/articles/dataset/TCGA_PanCanAtlas_Copy_Number_Data/6144122. We removed hypermutated samples (defined as five or more standard deviations above the mean non-silent somatic mutation count) from our dataset to reduce the number of false positives (i.e., non-driver mutations). After this filtering, 9,074 TCGA samples with mutation and copy number data remained. Any sample with a non-silent somatic variant in the target gene was included in the positive set. We also included copy number gains in the target gene for oncogenes and copy number losses in the target gene for tumor suppressor genes in the positive set; all remaining samples were considered negative for mutation in the target gene.

### Omics data download and preprocessing

RNA sequencing, 27K and 450K methylation array, microRNA, and RPPA datasets for TCGA samples were all downloaded from GDC, at the same link provided above. Mutational signatures information for TCGA samples with whole-exome sequencing data was downloaded from the International Cancer Genome Consortium (ICGC) data portal, at https://dcc.icgc.org/releases/PCAWG/mutational_signatures/Signatures_in_Samples/SP_Signatures_in_Samples. For our experiments, we used only the “single base signature” (SBS) mutational signatures, generated in [26]. In general, before training classifiers or extracting PCA components from all of the data types, we standardized (took z-scores of) each column/feature of all data types. For the RNA sequencing dataset, we generally used only the top 8,000 gene features by mean absolute deviation as predictors in our single-omics models, except where specified otherwise. For the RPPA, microRNA, and mutational signatures datasets, all columns/features were used.

To remove missing values from the methylation datasets, we removed the 10 samples with the most missing values, then performed mean imputation for probes with 1 or 2 values missing. All probes with missing values remaining after sample filtering and imputation were dropped from the analysis. This left us with 20,040 CpG probes in the 27K methylation dataset and 370,961 CpG probes in the 450K methylation dataset. For experiments where “raw” methylation data was used, we used the top 100,000 probes in the 450K dataset by mean absolute deviation for computational eiciency, and we used all of the 20,040 probes in the 27K dataset. For experiments where “compressed” methylation data was used, we used principal component analysis (PCA), as implemented in the `scikit-learn` Python library [36], to extract the top 5,000 principal components from the methylation datasets. We initially applied the beta-mixture quantile normalization (BMIQ) method [37] to correct for variability in signal intensity between type I and type II probes, but we observed that this had no effect on our results. We report uncorrected results in the main paper for simplicity.

### Construction of a set of cancer genes

To get a comprehensive picture of classification performance across a wide variety of cancer-related genes, we integrated several curated gene sets from the literature into a single “merged” cancer gene set. The individual gene sets we integrated were from Vogelstein et al. [38] (all genes from Table S2A), Bailey et al. [39] (only genes annotated as “pan-cancer” drivers in Table S1), and the COSMIC Cancer Gene Census [40] (all Tier 1 genes annotated as “somatic”). In addition, the COSMIC CGC dataset contains 3 possible “roles in cancer” for each gene: oncogene, TSG, and fusion gene; for this analysis we dropped genes that are annotated only as fusion genes (i.e. no oncogene or TSG annotation). These filters resulted in a starting dataset of 511 cancer-related genes, which we reduced further for each experiment based on the number of mutated (i.e. positively labeled) samples as described in the next section.

### Comparing data modalities

We made three main comparisons in this study: one between different sets of genes using only expression data, one comparing expression and DNA methylation data types, and one comparing all data types. This choice in comparisons was mainly due to sample size limitations, as running a single comparison using all data types would force us to use only samples that are profiled for every data type, which would discard a large number of samples that lack profiling on only one or a few data types. Thus, for each of the three comparisons, we used the intersection of TCGA samples having measurements for all of the datasets being compared in that experiment. This resulted in three distinct sets of samples: 9,074 samples shared between {expression, mutation} data, 7,981 samples shared between {expression, mutation, 27K methylation, 450K methylation}, and 5,226 samples shared between {expression, mutation, 27K methylation, 450K methylation, RPPA, microRNA, mutational signatures}. When we dropped samples between experiments as progressively more data types were added, we observed that the dropped samples had approximately the same cancer type proportions as the dataset as a whole. In other words, samples that were profiled for one data type but not another did not tend to come exclusively from one or a few cancer types. Exceptions included acute myeloid leukemia (LAML) which had no samples profiled in the RPPA data, and ovarian cancer (OV) which had only 8 samples with 450K methylation data. More detailed information on cancer type proportions profiled for each data type is provided in Supplementary Figure 8 and Supplementary Table 1.

For each target gene, in order to ensure that the training dataset was reasonably balanced (i.e. that there would be enough mutated samples to train an effective classifier), we included only cancer types with at least 15 mutated samples and at least 5% mutated samples, which we refer to here as “valid” cancer types. After applying these filters, the number of valid cancer types remaining for each gene varied based on the set of samples used: more data types resulted in fewer shared samples, and fewer samples generally meant fewer valid cancer types. In some cases, this resulted in genes with no valid cancer types, which we dropped from the analysis. Out of the 511 genes from the “merged” cancer gene set described in the previous section, for the analysis using {expression, mutation} data we retained 268 target genes, for the {expression, mutation, 27k methylation, 450k methylation} analysis we retained 272 genes, and for the analysis using all data types we retained 217 genes.

We additionally explored mutation prediction from gene expression alone using three gene sets of equal size: the cancer-related genes from the merged dataset described previously, a set of frequently mutated genes in TCGA, and a set of random genes with mutations profiled by MC3. To match the size of the merged cancer gene set, we took the 268 most frequently mutated genes in TCGA as quantified by MC3, all of which had at least one valid cancer type. For the random gene set, we first filtered to the set of all genes with one or more valid cancer types by the same criteria (15 total samples mutated and at least 5% of samples mutated), then sampled 268 of the resulting 1,348 genes uniformly at random. Based on the results of the gene expression experiments, we used the merged cancer-related gene set for all subsequent experiments comparing -omics data types.

### Training classifiers to detect cancer mutations

We trained logistic regression classifiers to predict whether or not a given sample had a mutational event in a given target gene using data from various -omics datasets as explanatory variables. Our model was trained on -omics data (*X*) to predict mutation presence or absence (*y*) in a target gene. To control for varying mutation burden per sample and to adjust for potential cancer type-specific expression patterns, we included one-hot encoded cancer type and log_10_(sample mutation count) in the model as covariates. Since our -omics datasets tend to have many dimensions and comparatively few samples, we used an elastic net penalty to prevent overfitting [41] in line with the approach used in Way et al. 2018 [9] and Way et al. 2020 [13]. Elastic net logistic regression finds the feature weights 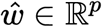 solving the following optimization problem:

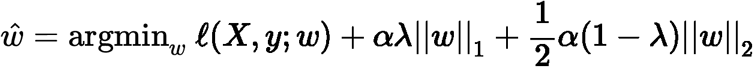

where *i* ∈ {1, …, *n*} denotes a sample in the dataset, *X*_*i*_ ∈ ℝ^*p*^ denotes features (omics measurements) from the given sample, *y*_*i*_ ∈ {0, 1} denotes the label (mutation presence/absence) for the given sample, and 𝓁(·) denotes the negative log-likelihood of the observed data given a particular choice of feature weights, i.e.

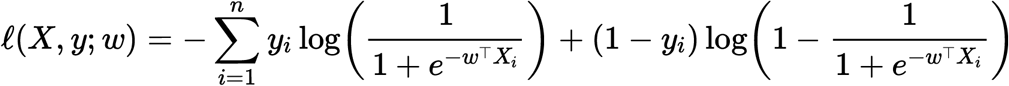

This optimization problem leaves two hyperparameters to select: *α* (controlling the tradeoff between the data log-likelihood and the penalty on large feature weight values), and *λ* (controlling the tradeoff between the L1 penalty and L2 penalty on the weight values). Although the elastic net optimization problem does not have a closed form solution, the loss function is convex, and iterative optimization algorithms are commonly used for finding reasonable solutions. For fixed values of *α* and *λ*, we solved for 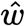 using stochastic gradient descent, as implemented in `scikit-learn` ‘s `SGDClassifier` method.

Given weight values 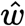, it is straightforward to predict the probability of a positive label (mutation in the target gene) 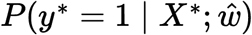 for a test sample *X*^∗^:

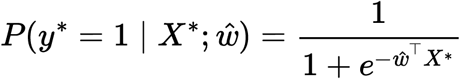

and the probability of no mutation in the target gene, 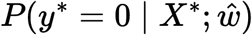, is given by (1 - the above quantity).

For each target gene, we evaluated model performance using two replicates of 4-fold cross-validation, where train and test splits were stratified by cancer type and sample type. That is, each training set/test set combination had equal proportions of each cancer type (BRCA, SKCM, COAD, etc.) and each sample type (primary tumor, recurrent tumor, etc.). To choose the elastic net hyperparameters, we used 3-fold nested cross-validation, with a grid search over the following hyperparameter ranges: *λ* = [0.0, 0.05, 0.1, 0.3, 0.5, 0.7, 0.9, 1.0] and *α* = [0.0001, 0.001, 0.01, 0.1, 1, 10]. Using the grid search results, for each evaluation fold we selected the set of hyperparameters with the optimal area under the precision-recall curve (AUPR), averaged over the three inner folds.

### Evaluating mutation prediction classifiers

Area under the receiver-operator curve (AUROC) [42] and area under the precision-recall curve (AUPR) [43] are metrics that are frequently used to quantify classification performance for a continuous or probabilistic output, such as that provided by logistic regression. These metrics summarize performance across a variety of binary label thresholds, rather than requiring choice of a single threshold to determine positive or negative predictions. In the main text, we report results using AUPR, summarized using average precision. AUPR has been shown to distinguish between models more accurately than AUROC when there are few positively labeled samples [44,45]. As an additional correction for imbalanced labels, in many of the results in the main text we report the difference in AUPR between a classifier fit to true mutation labels and a classifier fit to data where the mutation labels are randomly permuted. In cases where mutation labels are highly imbalanced (very few mutated samples and many non-mutated samples), a classifier with permuted labels may perform well simply by chance, e.g. by predicting the negative/non-mutated class for most samples. To maintain the same label balance for the classifiers with permuted labels as the classifiers with the true labels, we permuted labels separately in the train and test sets for each cross-validation split. Additionally, to maintain the same label proportions within each cancer type after permuting the labels, we permuted labels independently for each cancer type.

Recall that for each target gene and each -omics dataset, we ran two replicates of 4-fold cross-validation, for a total of eight performance results. To make a statistical comparison between two models using these performance distributions, we used paired-sample *t*-tests, where performance measurements derived from the same cross-validation fold are considered paired measurements. We used this approach to compare a model trained on true labels with a model trained on permuted labels (addressing the question, “for the given gene using the given data type, can we predict mutation status better than random”), and to compare a model trained on data type A with a model trained on data type B (addressing the question, “for the given gene, can we make more effective mutation status predictions using data type A or data type B”).

We corrected for multiple tests using a Benjamini-Hochberg false discovery rate correction. For experiments where we chose a binary threshold for accepting/rejecting *H*_0_ we set a conservative corrected threshold of *p* = 0.001; we were able to estimate the number of false positives by examining genes with better performance for permuted mutation labels than true labels. We chose this threshold to ensure that none of the observed false positive genes were considered significant, since we would never expect permuting labels to improve performance. However, our results were not sensitive to the choice of this threshold, and we display cutoffs of *p* = 0.05 and *p* = 0.01 in many of our plots as well.

### Survival prediction using -omics datasets

As a complementary comparison to mutation prediction, we constructed predictors of patient survival using the clinical data available from the GDC, in the `TCGA-CDR-SupplementalTableS1.xlsx` file. Following the methods described in [46], as the clinical endpoint we used overall survival (OS), except in nine cancer types with few deaths observed where we used progression-free intervals (PFI) as the clinical endpoint (BRCA, DLBC, LGG, PCPG, PRAD, READ, TGCT, THCA and THYM). For prediction, we used Cox regression as implemented in the Python package [47], with patient age at diagnosis and log_10_(sample mutation count) included as covariates, as well as a one-hot encoded variable for cancer type in the pan-cancer case. To ensure that the per-feature information content was comparable between -omics data types, we preprocessed the -omics datasets using PCA and extracted the top *k* principal components; in the case where the number of features in the original dataset was less than *k* we used all available PCs (that is, we set *k* = min(*p, k*) where *p* is the number of features in the unprocessed dataset). For the pan-cancer models we plot results over multiple values of *k*: *k* ∈ {10, 100, 500, 1000, 5000}; for the individual cancer type models we set *k* = 10.

To model pan-cancer survival (results shown in main paper), we used the elastic net Cox regression implementation in `scikit-survival` (i.e. the `CoxnetSurvivalAnalysis` method). To select hyperparameters for the elastic net Cox regression model, we performed a grid search over *λ* = [0.0, 0.05, 0.1, 0.3, 0.5, 0.7, 0.9, 1.0] and *α* = [0, 1e-5, 1e-4, 5e-4, 0.001, 0.005, 0.01, 0.05, 0.1, 0.5, 1, 10, 100, 1000]. To model survival using data from individual cancer types (results shown in supplement), we found elastic net Cox regression to be unstable even for small feature dimensions, so we used the ridge-regularized Cox regresssion implementation in the `CoxPHSurvivalAnalysis` method in `scikit-survival`. To select the regularization parameter *α*, we used the default selection procedure implemented in `scikit-survival` to determine a range of potential *α* values based on the data. This procedure begins by deriving the maximum *α* value as the smallest value for which all coeicients are 0 (call this *α*_max_), then it selects 100 possibilities for alpha spaced evenly on a log scale between *α*_max_ and 0.01 · *α*_max_. We found that for individual cancer types, this data-driven procedure resulted in more consistent and stable model convergence than choosing a fixed set of alphas to search over as in the pan-cancer survival prediction experiments.

We measured survival prediction performance using the censored concordance index (c-index) [48], which quantifies agreement between the order of survival time predictions and true outcomes for a held-out dataset; higher c-index values indicate more accurate survival prediction performance. Similar to the mutation prediction experiments, we calculated c-index values on held-out subsets of the data for two replicates of 4-fold cross-validation, resulting in eight performance measurements for each model. As a baseline, for both the pan-cancer and cancer type specific datasets, we constructed survival models using only non-omics covariates. For the pan-cancer data, covariates included patient age at diagnosis, log_10_(sample mutation count), and a one-hot encoded variable for sample cancer type. The cancer type-specific baseline models were the same, without the cancer type indicator, since all train and test samples were derived from the same cancer type.

### Multi-omics mutation prediction experiments

To predict mutation presence or absence in cancer genes using multiple data types simultaneously, we concatenated individual datasets into a large feature matrix, then used the same elastic net logistic regression method described previously. For this task, we considered only the gene expression, 27K methylation, and 450K methylation datasets. We used only these data types to limit the number of multi-omics combinations; the expression and methylation datasets resulted in the best overall performance across the single-omics experiments, so we limited combinations to those datasets. In the main text, we report results using the top 5,000 principal components for each dataset, which ensures that most variance is captured (approximately 95-98% of variance for each data type). In the supplement, we also report results using “raw” features: for gene expression we used all 15,639 genes available in our RNA sequencing dataset, and for the 27K and 450K methylation datasets we used the top 20,000 CpG probes by mean absolute deviation.

To construct the multi-omics models, we considered each of the pairwise combinations of the datasets listed above, as well as a combination of all 3 datasets. When combining multiple datasets, we concatenated along the column axis and included covariates for cancer type and sample mutation burden as before. For all multi-omics experiments, we used only the samples from TCGA with data for all three data types (i.e. the same 7,981 samples used in the single-omics experiments comparing expression and methylation data types). We considered a limited subset of well-performing genes from the merged cancer gene set as target genes, including *EGFR, IDH1, KRAS, PIK3CA, SETD2*, and *TP53*. We selected these genes because we had previously observed that they have good predictive performance and because they represent a combination of alterations that have strong gene expression signatures (*KRAS, EGFR, IDH1, TP53*) and strong DNA methylation signatures (*IDH1, SETD2, TP53*).

For the experiments predicting mutation status using a 3-layer fully connected neural network, described in the Results section and the supplement, we used the top 5,000 principal components as input for each data type. We selected hyperparameters for each of the 8 outer cross-validation splits using a single inner train/validation split and a random search over 20 hyperparameter combinations. The hyperparameter ranges that we sampled from in the random search were: `learning_rate: [0.1, 0.01, 0.001, 5e-4, 1e-4], h1_size: [1000, 500, 250], dropout: [0.5, 0.75, 0.9], weight_decay: [0, 0.1, 1, 100]`. Here, `h1_size` refers to the size of the first hidden layer, and the size of the second hidden layer was always set to `h1_size / 2`. As in the elastic net grid search, we chose the combination of hyperparameters with the best AUPR on the validation set, and retrained the model with those hyperparameters to make predictions on the test set. We trained our networks with the Adam optimizer [49], with ReLU activation after the hidden layers and sigmoid activation to make predictions, and using binary cross-entropy as the loss function as implemented in the PyTorch `BCEWithLogitsLoss` function, through the `skorch` library which provides interoperability between PyTorch and scikit-learn.

## Data and code availability

All analyses were implemented in the Python programming language and are available in the following GitHub repository: https://github.com/greenelab/mpmp under the open-source BSD 3-clause license. Scripts to download large data files from GDC and other sources are located in the `00_download_data` directory. Scripts to run experiments comparing data modalities used individually are located in the `02_classify_mutations` directory, scripts to run multi-omics experiments are located in the `05_classify_mutations_multimodal` directory, and scripts to run survival prediction experiments are located in the `06_predict_survival` directory. The Python environment was managed using `conda`, and directions for setting up the environment can be found in the `README.md` file. Most analyses were run on the HTC CPU cluster at the University of Pittsburgh, except the neural networks which were trained and evaluated on the PMACS LPC GPU cluster at the University of Pennsylvania; scripts for training classifiers both locally for a single gene and on a Slurm cluster to reproduce the analysis of many genes in parallel are provided in the linked GitHub repo. This manuscript was written using Manubot [50] and is available on GitHub at https://github.com/greenelab/mpmp-manuscript.

## Results

### Using diverse data modalities to predict cancer alterations

We collected five different data modalities from cancer samples in the TCGA Pan-Cancer Atlas, capturing five steps of cellular function that are perturbed by genetic alterations in cancer (Figure 1A). These included gene expression (RNA-seq data), DNA methylation (27K and 450K Illumina BeadChip arrays), protein abundance (RPPA data), microRNA expression data, and patterns of somatic mutation (mutational signatures). To link these diverse data modalities to changes in mutation status, we used elastic net logistic regression to predict the presence or absence of mutations in cancer genes, using these readouts as predictive features (Figure 1B). We evaluated the resulting mutation status classifiers in a pan-cancer setting, preserving the proportions of each of the 33 cancer types in TCGA for eight train/test splits (4 folds x 2 replicates) in each of approximately 250 cancer genes (Figure 1C).

**Figure 1:**
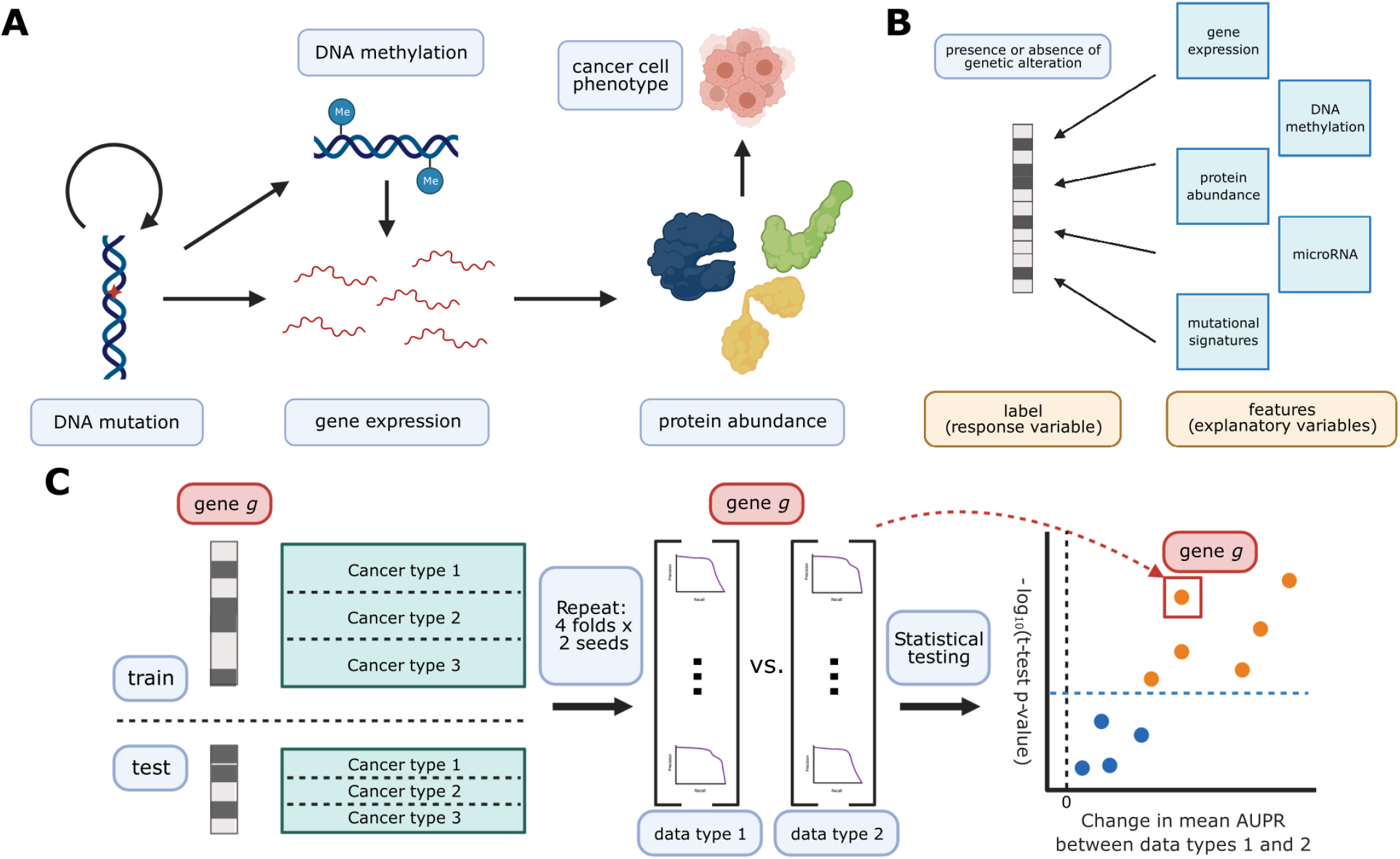
**A**. Cancer mutations can perturb cellular function via a variety of cellular processes. Arrows represent major potential paths of information flow from a somatic mutation in DNA to its resulting cell phenotype; circular arrow represents the ability of certain mutations (e.g. in DNA damage repair genes) to alter somatic mutation patterns. Note that this does not reflect all possible relationships between cellular processes: for instance, changes in gene expression can lead to changes in somatic mutation rates. **B**. Predicting presence/absence of somatic alterations in cancer from diverse data modalities. In this study, we use functional readouts from TCGA as predictive features and the presence or absence of mutation in a given gene as labels. This reverses the primary direction of information flow shown in Panel A. **C**. Schematic of evaluation pipeline.

We sought to compare classifiers against a baseline where mutation labels are permuted (to identify genes whose mutation status correlates strongly with a functional signature in a given data type) and also to compare classifiers trained on true labels across different data types (to identify data types that are more or less predictive of mutations in a given gene). To account for variation between dataset splits in making these comparisons, we treat classification metrics from the eight train/test splits as performance distributions, which we compare using *t*-tests. We summarize performance across all genes in our cancer gene set using a similar approach to a volcano plot, in which each point is a gene. In our summary plots, the x-axis shows the magnitude of the change in the classification metric between conditions, and the y-axis shows the *p*-value for the associated *t*-test (Figure 1C).

### Selection of cancer-related genes improves predictive signal

As a baseline, we evaluated prediction of mutation status from gene expression data across several different gene sets. Past work has evaluated mutation prediction for the top 50 most mutated genes in TCGA [13], and we sought to extend this to a broader list of gene sets. To evaluate whether using known cancer-related genes tends to improve prediction, we compiled a set of cancer-related genes (n=268) from Vogelstein et al. 2013 [38], Bailey et al. 2018 [39], and the COSMIC Cancer Gene Census [40]. We compared performance on this curated gene set with performance on an equal number of genes sampled randomly after applying a mutation frequency threshold (n=268, see Methods for sampling details) and an equal number of the most mutated genes in TCGA (n=268). For all gene sets, we used only the set of TCGA samples for which both gene expression and somatic mutation data exists, resulting in a total of 9,074 samples across all 33 cancer types. This set of samples was further filtered for each target gene to cancer types containing at least 15 mutated samples and at least 5% of samples mutated for that cancer type. As an alternate approach, we tried including/excluding entire genes using similar filters, and the results were consistent across filtering strategies (Supplementary Figure 11). We then evaluated the performance for each target gene in each of the three gene sets.

Overall, genes from the cancer-related gene set were more predictable than randomly chosen genes or those selected by total mutation count (Figure 2A). In total, for a significance threshold of *α* = 0.001, 120/268 genes (44.8%) in the cancer-related gene set are significantly predictable from gene expression data, compared to 14/268 genes (5.22%) in the random gene set and 80/268 genes (29.9%) in the most mutated gene set. Of the 14 significantly predictable genes in the random gene set, 13 of them are also in the cancer-related gene set (highlighted with ‘X’ in Figure 2B), and of the 80 significantly predictable genes in the most mutated gene set, 26 of them are also in the cancer-related gene set (highlighted in red in Figure 2C). These results suggest that selecting target genes for mutation prediction based on prior knowledge of their involvement in cancer pathways and processes, rather than randomly or based on mutation frequency alone, can improve predictive signal and identify more highly predictable mutations from gene expression data.

**Figure 2:**
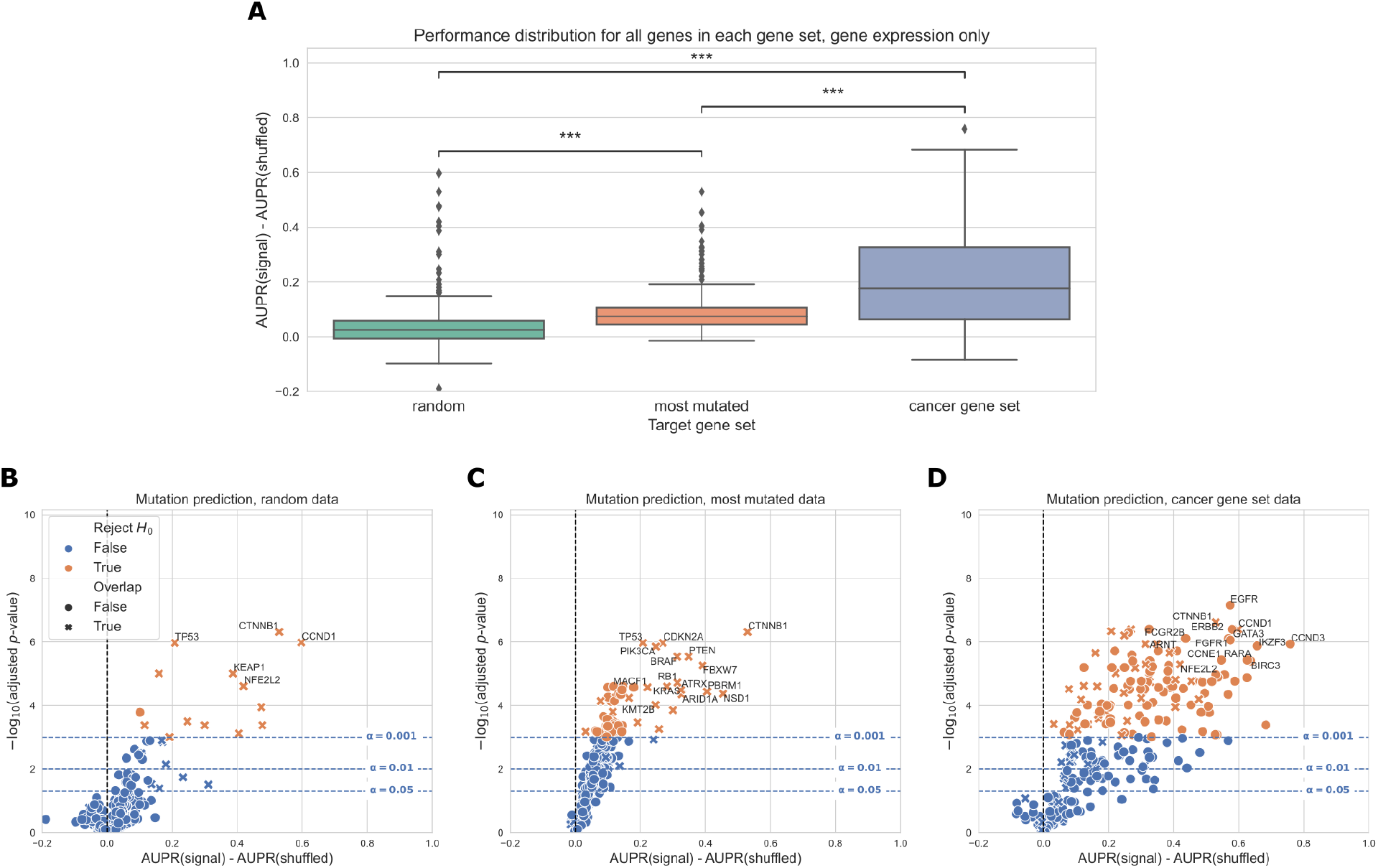
**A**. Overall distribution of performance across three gene sets, using gene expression (RNA-seq) data to predict mutations. Each data point represents the mean cross-validated AUPR difference, compared with a baseline model trained on permuted mutation presence/absence labels, for one gene in the given gene set; notches show bootstrapped 95% confidence intervals. “random” = 268 random genes, “most mutated” = 268 most mutated genes, “cancer gene set” = 268 cancer related genes from curated gene sets. Significance stars indicate results of Bonferroni-corrected pairwise Wilcoxon tests: **: *p* < 0.01, ***: *p* < 0.001, ns: not statistically significant for a cutoff of *p* = 0.05. **B, C, D**. Volcano-like plots showing mutation presence/absence predictive performance for each gene in each of the three gene sets. The *x*-axis shows the difference in mean AUPR compared with a baseline model trained on permuted labels, and the *y*-axis shows *p*-values for a paired *t*-test comparing cross-validated AUPR values within folds. Points (genes) marked with an “X” are overlapping between the cancer gene set and either the random or most mutated gene set.

### Gene expression predicts cancer mutation status more effectively than DNA methylation

We compared gene expression with DNA methylation as downstream readouts of the effects of cancer alterations. In these analyses, we considered both the 27K probe and 450K probe methylation datasets generated for the TCGA Pan-Cancer Atlas. As target genes, we used the same combined cancer-related gene set described in the “Selection of cancer-related genes” section. We used samples that had data for each of the data types being compared, including somatic mutation data to generate mutation labels. This process retained 7,981 samples in the intersection of the expression, 27K methylation, 450K methylation, and mutation datasets, which we used for subsequent analyses. The most frequent missing data types were somatic mutation data (1,114 samples) and 450K methylation data (1,072 samples) (Figure 3A).

**Figure 3:**
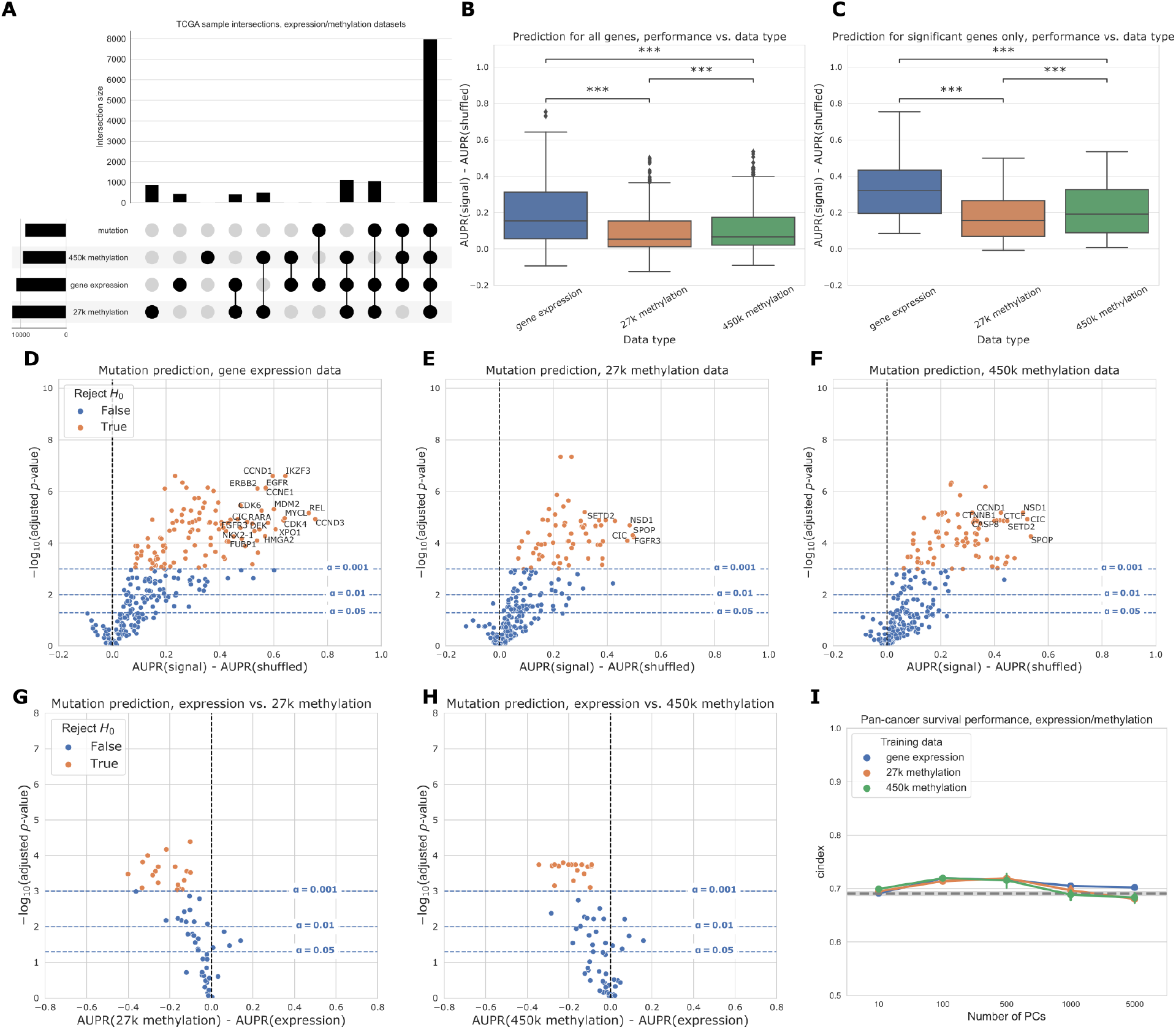
**A**. Count of overlapping samples between gene expression, 27K methylation, 450K methylation, and somatic mutation data used from TCGA. Only non-zero overlap counts are shown. Somatic mutation sample information is included because it is needed to generate the mutation presence/absence labels. **B**. Predictive performance for genes in the cancer-related gene set, using each of the three data types as predictors. The gene expression predictor uses the top 8000 gene features by mean absolute deviation, and the methylation predictors use the top 5000 principal components as predictive features. Significance stars indicate results of Bonferroni-corrected pairwise Wilcoxon tests: **: *p* < 0.01, ***: *p* < 0.001, ns: not statistically significant for a cutoff of *p* = 0.05. **C**. Predictive performance for genes where at least one of the considered data types predicts mutation labels significantly better than the permuted baseline. **D-F**. Predictive performance for each gene in the cancer-related gene set, for each data type, compared with a baseline model trained on permuted labels. **G-H**. Direct comparison of performance using gene expression and each methylation dataset, for genes that perform significantly better than the baseline for both data types. Points (genes) to the left of y=0 perform better using gene expression-derived features, and points to the right perform better using methylation-derived features. **I**. Pan-cancer survival prediction performance, quantified using c-index on the *y*-axis, for gene expression, 27K methylation, and 450K methylation. The *x*-axis shows results with varying numbers of principal components included for each data type. Models also included covariates for patient age, sample mutation burden, and sample cancer type; grey dotted line indicates mean performance for a covariate-only baseline model.

For many genes, predictions are better than our baseline model where labels are permuted (values greater than 0 in the box plots), suggesting that there is considerable predictive signal in both expression and methylation datasets across the cancer-related gene set (Figure 3B). On aggregate across all genes, predictive performance is best overall for gene expression. Both before and after filtering for genes that exceed the significance threshold, gene expression with raw gene features provides a significant performance improvement relative to the 27K methylation and 450K methylation datasets (Figure 3B-C). Results were similar with PCA-compressed gene expression features or raw CpG probes as predictors (Supplementary Figure 12).

Considering each target gene in the cancer-related gene set individually, we observed that 113/272 genes significantly outperformed the permuted baseline using gene expression data, as compared to 62/272 genes for 27K methylation and 77/272 genes for 450K methylation (Figure 3D-F, more information about specific genes in Supplementary Figure 9). Some “well-predicted” genes that outperformed the permuted baseline tended to be similar between data types (Figure 3D-F; genes in the top right of each plot). For example, *CIC* appears in the top right of all 3 plots, and *CCND1* appears in the top right of the gene expression and 450K methylation plots, suggesting that mutations in these genes have strong gene expression and DNA methylation signatures, and these signatures tend to be preserved across cancer types.

In addition to comparing mutation classifiers trained on different data types to the permuted baseline, we also compared classifiers trained on true labels directly to each other, for genes that performed significantly better than the baseline for both of the data types under consideration (Figure 3G-H). We observed that 18/58 genes were significantly more predictable from expression data than 27K methylation data, and 21/69 genes were significantly more predictable from expression data than 450K methylation data. In both cases, no genes were significantly more predictable using the methylation data types. Still, we observed that some points were clustered around the origin, indicating that the data types appear to confer similar information about mutation status. That is, in these cases, matching the gene being studied with the “correct” data modality seems to be unimportant: mutation status has a strong signature which can be extracted from both expression and DNA methylation data roughly equally.

We additionally compared pan-cancer survival prediction performance using principal components derived from each data type; in general, results were comparable across the three data types (Figure 3I). All data types outperformed the covariate-only baseline (see Methods) for lower numbers of PC features included, although performance was similar to the baseline for higher numbers of PCs. Confidence intervals between the best- and worst-performing data types overlap at most PC counts (with the exception of gene expression at 5,000 PC features), suggesting that similarly to mutation prediction, the three data types tend to have comparable effectiveness for pan-cancer survival prediction.

Focusing on several selected genes of interest, we observed that relative classifier performance varies by gene (Figure 4). Past work has indicated that mutations in *TP53* are highly predictable from gene expression data [11], and we observed that the methylation datasets provided similar predictive performance (Figure 4A). Similarly, for *IDH1* both expression and methylation features result in similar performance, consistent with the previously observed role of IDH1 in regulating both DNA methylation and gene expression (Figure 4D) [51]. Mutations in *KRAS* and *ERBB2* (*HER2*) were most predictable from gene expression data, and in both cases the methylation datasets significantly outperformed the baseline as well (Figure 4B and 4E). Gene expression signatures of *ERBB2* alterations are historically well-studied in breast cancer [52], and samples with activating *ERBB2* mutations have recently been shown to share sensitivities to some small-molecule inhibitors across cancer types [53]. These observations are consistent with the pan-cancer *ERBB2* mutant-associated expression signature that we observed in this study. *NF1* mutations were also most predictable from gene expression data, although the gene expression-based *NF1* mutation classifier did not significantly outperform the baseline with permuted labels at a cutoff of *α* = 0.001 (Figure 4C). *SETD2* is an example of a gene that is more predictable from the methylation datasets than from gene expression, although gene expression with raw gene features significantly outperformed the permuted baseline as well (Figure 4F). *SETD2* is widely mutated across cancer types and affects H3K36 histone methylation most directly, but SETD2-mediated changes in H3K36 methylation have been linked to dysregulation of diverse cellular processes including DNA methylation and RNA splicing [23,54].

**Figure 4:**
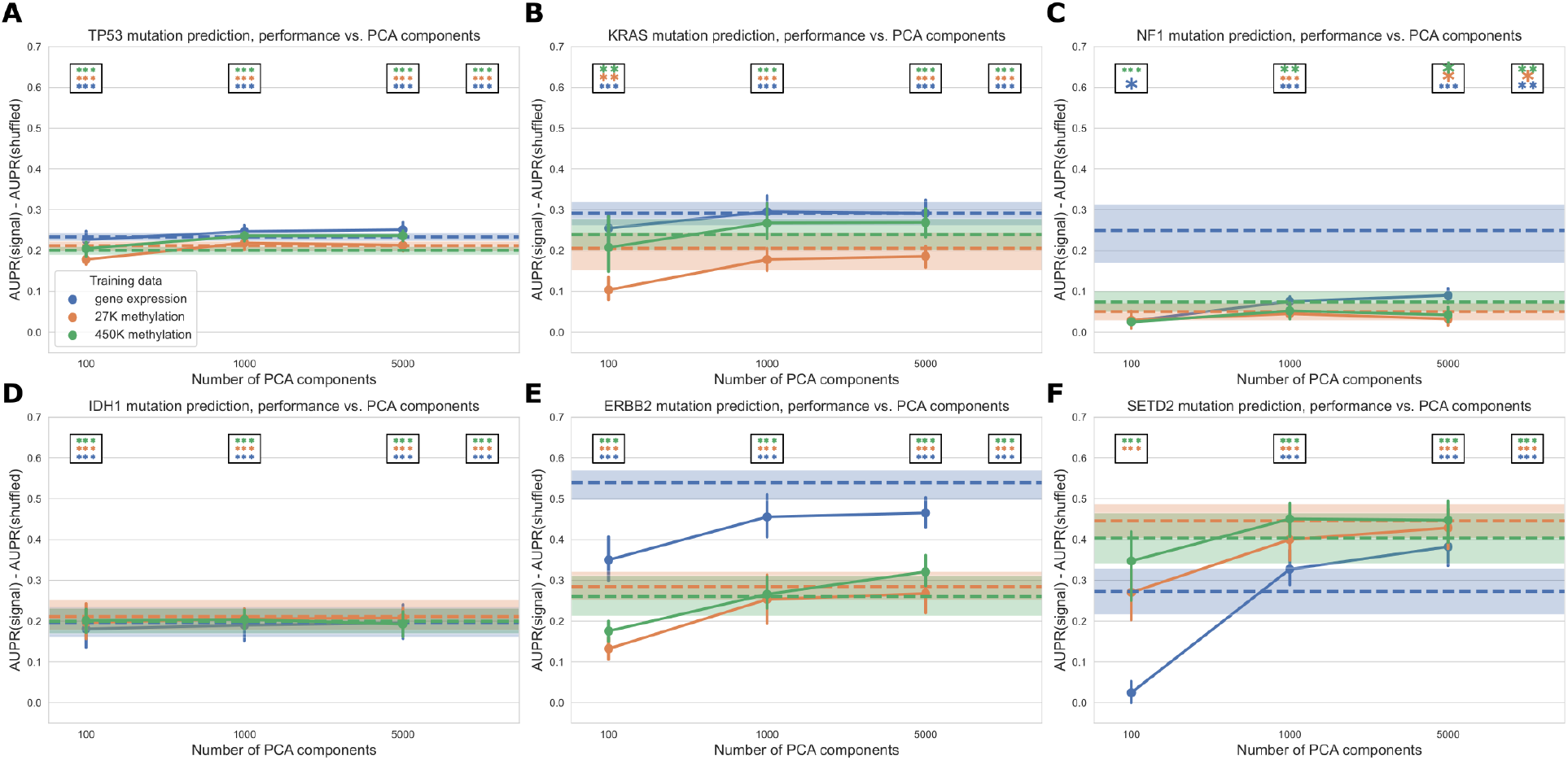
Performance across varying PCA dimensions for specific genes of interest. Dotted lines represent results for “raw” features (8,000 gene features for gene expression data and 8,000 CpG probes for both methylation datasets, selected by largest mean absolute deviation). Error bars and shaded regions show bootstrapped 95% confidence intervals. Stars in boxes show statistical testing results compared with permuted baseline model; each box refers to the model using the number of PCA components it is over (far right box = models with raw features). **: *p* < 0.01, ***: *p* < 0.001, no stars: not statistically significant for a cutoff of *p* = 0.05.

### Comparing six different readouts favors expression and DNA methylation

Next, we expanded our comparison to all five functional data modalities (six total readouts, since there are two DNA methylation platforms) available in the TCGA Pan-Cancer Atlas. As with previous experiments, we limited our comparison to the set of samples profiled for each readout, resulting in 5,226 samples with data for all readouts. The data types with the most missing samples were RPPA data (2,215 samples that were missing RPPA data) and 450K methylation (630 samples that were missing 450K methylation data) (Figure 5A). Summarized over all genes in the cancer-related gene set, we observed that gene expression tended to produce better predictions than the other data types (Figure 5B). This remained true when we looked only at the set of genes having at least one significant predictor (i.e. “well-predicted” genes) (Figure 5C).

**Figure 5:**
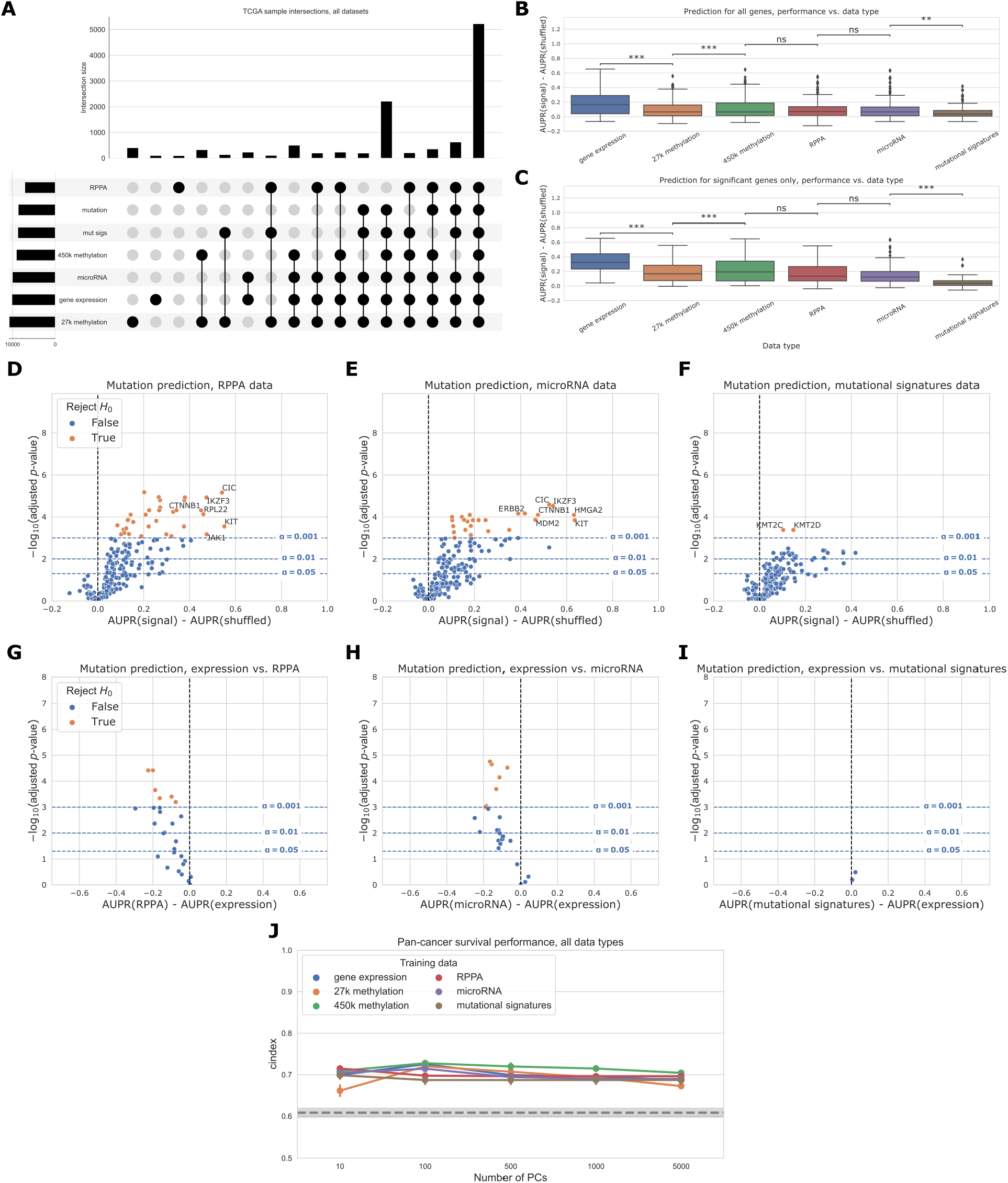
**A**. Overlap of TCGA samples between all data types used in mutation prediction comparisons. Only overlaps with more than 100 samples are shown. Somatic mutation sample information is included because it is needed to generate the mutation presence/absence labels. **B**. Overall distribution of performance per data type across 217 genes from the cancer-related gene set. Each data point represents mean cross-validated AUPR difference, compared with a baseline model trained on permuted labels, for one gene; notches show bootstrapped 95% confidence intervals. Significance stars indicate results of Bonferroni-corrected pairwise Wilcoxon tests: **: *p* < 0.01, ***: *p* < 0.001, ns: not statistically significant for a cutoff of *p* = 0.05. All pairwise tests were run, and corrected for, but only neighboring test results are shown. **C**. Overall performance distribution per data type for genes where the permuted baseline model is significantly outperformed for one or more data types, resulting in a total of 39 genes. **D, E, F**. Volcano-like plots showing predictive performance for each gene in the cancer-related gene set, in each of the added data types (RPPA, microRNA, mutational signatures). The *x*-axis shows the difference in mean AUPR compared with a baseline model trained on permuted labels, and the *y*-axis shows *p*-values for a paired *t*-test comparing cross-validated AUPR values within folds. **G, H, I**. Direct comparison of performance using gene expression and each added data type, showing only genes that perform significantly better than the baseline model for both data types. Points (genes) to the left of y=0 perform better using gene expression-derived features, and points to the right perform better using the added data type (RPPA, microRNA, and mutational signatures respectively). **J**. Pan-cancer survival prediction performance, quantified using c-index on the *y*-axis, for all data types. The *x*-axis shows results with varying numbers of principal components included for each data type. Models also included covariates for patient age, sample mutation burden, and sample cancer type; grey dotted line indicates mean performance for a covariate-only baseline model.

On the individual gene level, mutations in 33/217 genes were significantly predictable from RPPA data relative to the permuted baseline, compared to 25/217 genes from microRNA data and 2/217 genes from mutational signatures data (Figure 5D-F). For the remaining data types on this smaller set of samples, 79/217 genes outperformed the baseline for gene expression data, 31/217 for 27k methylation, and 42/217 for 450k methylation. Compared to the methylation experiments (Figure 3), we observed fewer “well-predicted” genes for the expression and methylation datasets here (likely due to the considerably smaller sample size) but relative performance was comparable (Supplementary Figure 10). Direct comparisons between each added data type and gene expression data showed that for most “well-predicted” genes, RPPA, microRNA and mutational signatures data generally provide similar or worse performance compared to gene expression (Figure 5G-I).

Performance using RPPA data (Figure 5G) is notable because of its drastically smaller dimensionality than the other data types (190 proteins, compared to thousands of features for the expression and methylation data types). This suggests that each protein abundance measurement provides a high information content, although this is by design as the antibody probes used for the TCGA analysis were selected to cover established cancer-related pathways [55]. Similarly, the scope of the features captured by the mutational signatures data we used is limited to single-base substitution signatures; a broader spectrum of possible signatures is described in previous work [26,56] including doublet-substitution signatures, small indel signatures, and signatures of structural variation, but these were not readily available for the TCGA exome sequencing data. The relatively poor predictive ability of mutational signatures likely stems from a combination of biological and technical factors, as there is no reason to expect that changes in somatic mutation patterns would be directly caused by most cancer driver mutations. Two exceptions are *KMT2C* and *KMT2D* (Figure 5F), which may have a role in mediating DNA damage response [57].

As in the expression/methylation comparison, we also compared pan-cancer survival prediction performance between all six readouts, using the top principal components derived from each data type to ensure comparable information content (Figure 5J). All six readouts performed comparably for this smaller set of samples, with slightly better performance across PC feature dimensions for the 450K methylation array. The covariate-only baseline predictor performed considerably worse than it did in the expression/methylation comparisons, with all -omics data types outperforming the baseline predictor at all PC numbers.

When we constructed a heatmap depicting predictive performance for each gene across data types, we found that many genes tended to be well-predicted by more than one data type (Figure 6). Of the 86 genes that are well-predicted using at least one data type (grey circles in Figure 6), 52/86 (60.5%) are well-predicted by multiple data types, meaning that multiple -omics readouts contain a detectable signature of presence/absence of a mutation in the corresponding gene. Of the remaining 34 genes, 28/34 (82.4%) are well-predicted by gene expression alone. This supports our observation that in a surprising number of cases, choosing the “correct” data modality is unimportant for driver genes with strong functional signatures, although gene expression may be the best “default” choice as it tends to be a strong predictor in the majority of cases. Exceptions included *ERBB4, KMT2A, PIK3R1*, and *RPL22* (only well-predicted using RPPA data), *FAT4* (only well-predicted using microRNA data), and *KDM6A* (only well-predicted using 450K methylation data).

**Figure 6:**
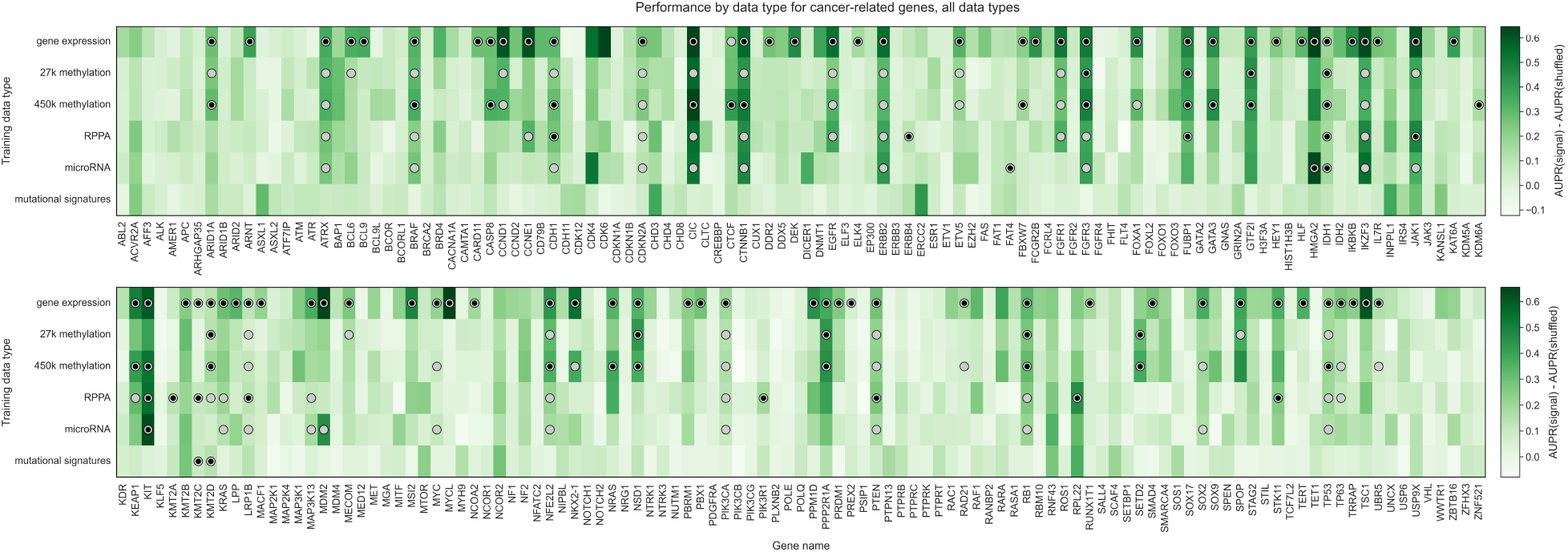
Heatmap displaying predictive performance for mutations in each of the 217 genes from the cancer-related gene set, across all six TCGA data modalities. Each cell quantifies performance for a target gene, using predictive features derived from a particular data type. Grey shaded dots indicate that the given data type provides significantly better predictions than the permuted baseline for the given gene; black inner dots indicate the same and additionally that the given data type provides statistically equivalent performance to the data type with the best average performance (determined by pairwise *t*-tests across data types with FDR correction).

### Simple multi-omics integration provides little performance benefit

We also trained “multi-omics” classifiers to predict mutations in six well-studied and widely mutated driver genes across various cancer types: *EGFR, IDH1, KRAS, PIK3CA, SETD2*, and *TP53*. Each of these genes is well-predicted from several data types in our earlier experiments (Figure 6), consistent with having strong pan-cancer driver effects. For the multi-omics classifiers, we considered all pairwise combinations of the top three performing individual data types (gene expression, 27K methylation, and 450K methylation), in addition to a model using all three data types. We trained a classifier for multiple data types by concatenating features from the individual data types, then fitting the same elastic net logistic regression model as we used for the single-omics models. Here, we show results using the top 5,000 principal components from each data type as predictive features, to ensure that feature count and scale is comparable among data types; results for raw features are shown in Supplementary Figure 13. We additionally ran the same experiments using a 3-layer fully-connected neural network for classification, with principal components as input, and results are shown in Supplementary Figure 14. In general, we found predictions using elastic net logistic regression to be more robust across cross-validation folds and hyperparameter choices than predictions using the neural network, although the neural network provided a slight performance improvement using multiple -omics types for some genes.

For each of the six target genes, we observed comparable performance between the best single-omics classifier (blue boxes in Figure 7A) and the best multi-omics classifier (orange boxes in Figure 7A). Across all classifiers and data types, we found varied patterns based on the target gene. For *IDH1* and *TP53* performance is relatively consistent regardless of data type(s), suggesting that baseline performance is high and there is little room for improvement as data is added (Figure 7C, G). The *TP53* classifier using raw features showed a statistically significant improvement when multiple data types were integrated, although the difference in mean performance was relatively small (Supplementary Figure 13, *p*=0.0078). For *EGFR, KRAS*, and *PIK3CA*, combining gene expression with methylation data results in statistically equivalent or worse performance to gene expression alone; classifiers trained only on methylation data generally do not perform as well as those that integrate expression data (Figure 7B, D, E). Previously, we saw that the best classifiers for *SETD2* used methylation data alone (Figure 6). When we added multiple data types to our *SETD2* classifier, we did observe an increase in performance (Figure 7F), although this improvement was not statistically significant in a paired-sample *t*-test for *α*=0.05 (*p*=0.078). Overall, we observed that combining data types in a relatively simple manner, by concatenating features from each individual data type, provided little or no improvement in predictive ability over the best individual data type. This supports our earlier observations of the redundancy of gene expression and methylation data as functional readouts, since our multi-omics classifiers are not in general able to extract gains in predictive performance as more data types are added for this set of cancer drivers.

**Figure 7:**
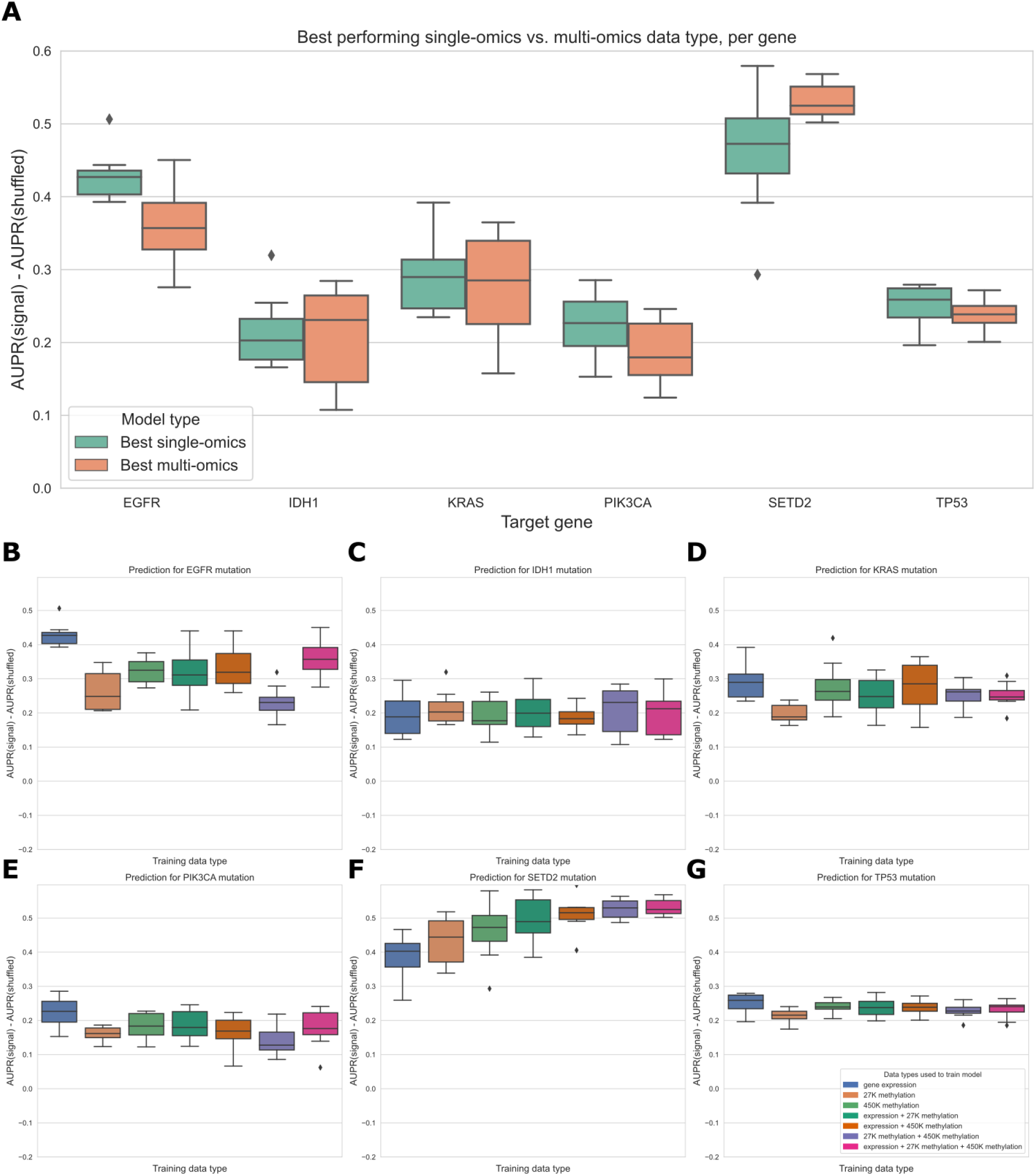
**A**. Comparing the best-performing model (i.e. highest mean AUPR relative to permuted baseline) trained on a single data type against the best “multi-omics” model for each target gene. None of the differences between single-omics and multi-omics models were statistically significant using paired-sample Wilcoxon tests across cross-validation folds, for a threshold of 0.05. **B-G**. Classifier performance, relative to baseline with permuted labels, for mutation prediction models trained on various combinations of data types. Each panel shows performance for one of the six target genes; box plots show performance distribution over 8 evaluation sets (4 cross-validation folds x 2 replicates).

**Figure 8:**
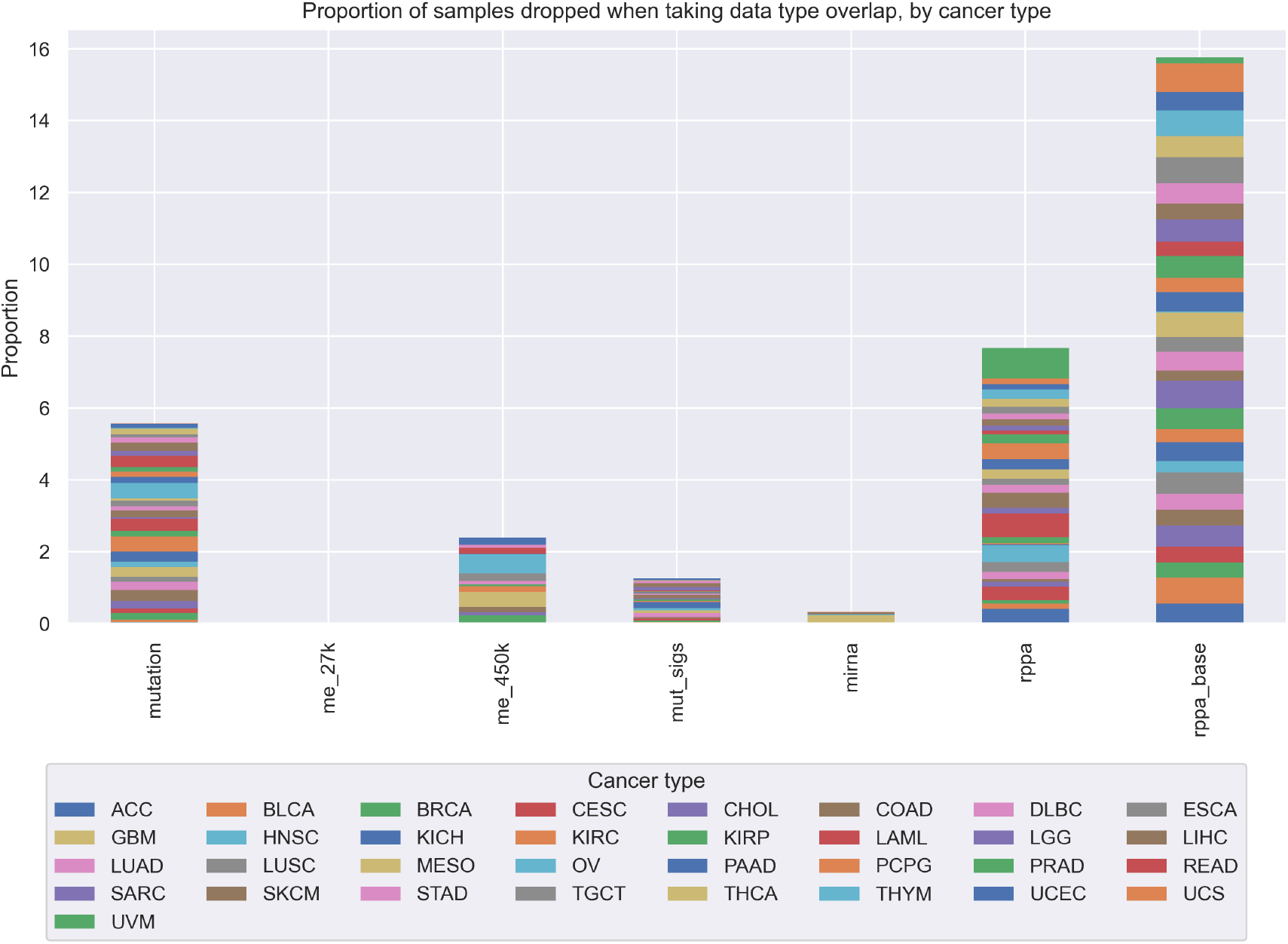
Proportion of samples from each TCGA cancer type that are “dropped” as more data types are added to our analyses. We started with gene expression data, and for each added data type, we took the intersection of samples that were profiled for that data type and the previous data types, dropping all samples that were missing 1 or more data types. Overall, at each step, the proportions of “dropped” samples appear to be fairly evenly spread between cancer types, showing that in general we are not disproportionately losing one or several cancer types as more data modalities are added to our analyses.

**Figure 9:**
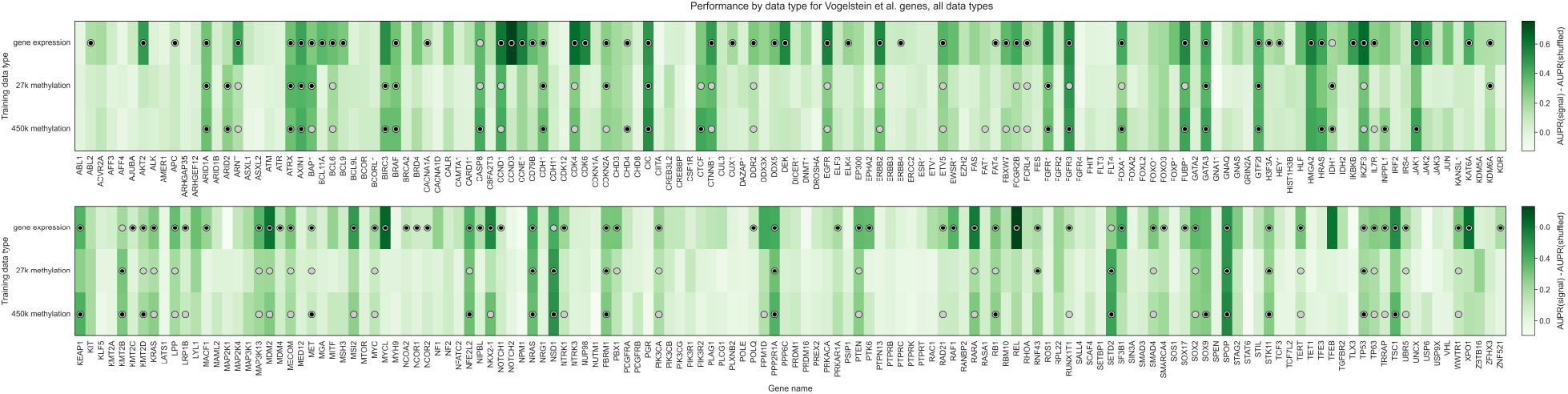
Heatmap displaying predictive performance for mutations in each of the 272 genes from the cancer-related gene set, across gene expression and the two DNA methylation arrays. Each cell quantifies performance for a target gene, using predictive features derived from a particular data type. Grey shaded dots indicate that the given data type provides significantly better predictions than the permuted baseline for the given gene; black inner dots indicate the same and additionally that the given data type provides statistically equivalent performance to the data type with the best average performance (determined by pairwise *t*-tests across data types with FDR correction).

**Figure 10:**
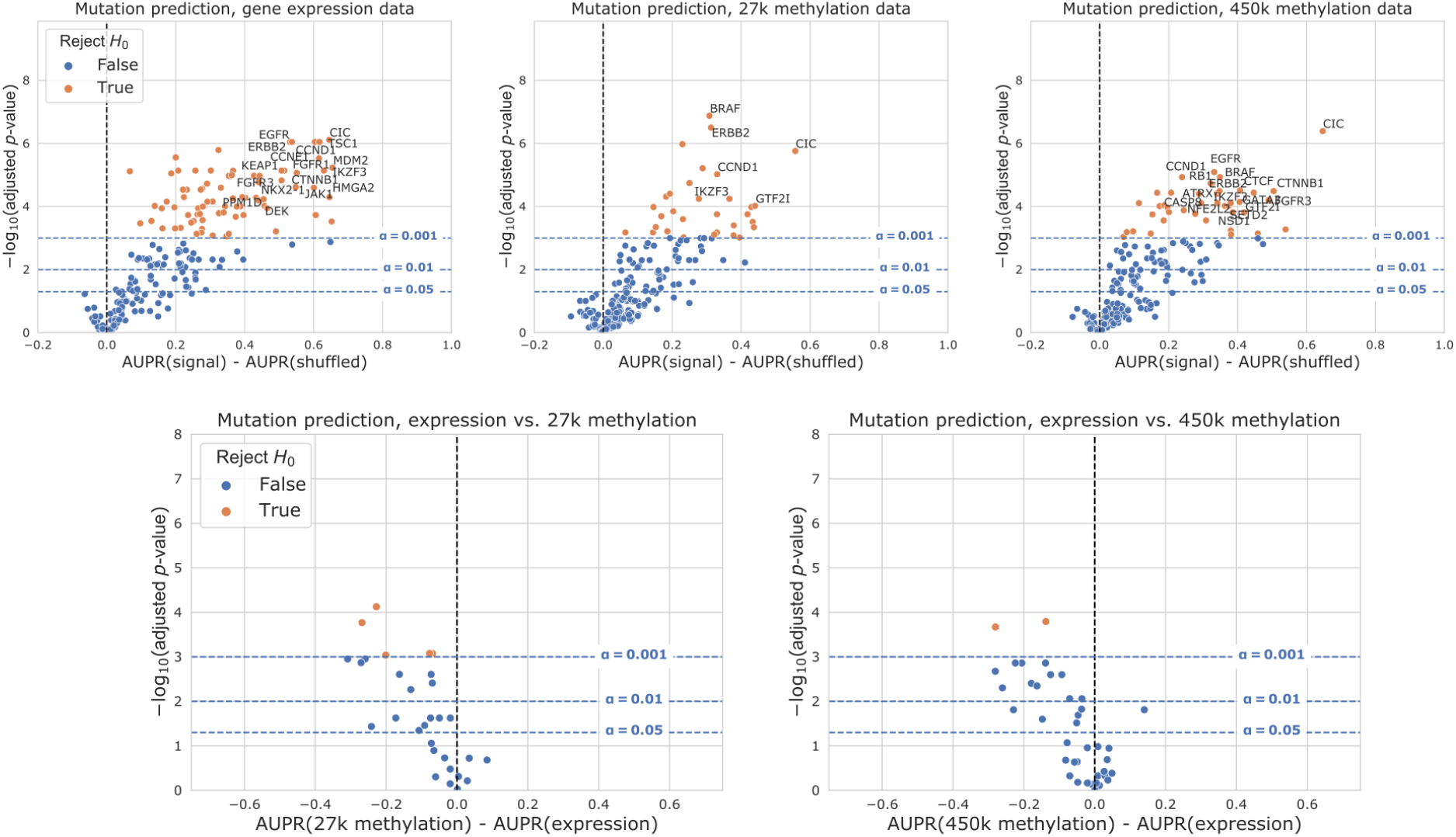
Volcano-like plots showing predictive performance for each gene in the cancer-related gene set for expression and DNA methylation, on the sample set used for the “all data types” experiments. The first row shows performance relative to the permuted baseline, and the second row shows direct comparisons between data types for genes that outperformed the permuted baseline only for both data types. The *x*-axis shows the difference in mean AUPR compared with a baseline model trained on permuted labels, and the *y*-axis shows *p*-values for a paired *t*-test comparing cross-validated AUPR values within folds.

**Figure 11:**
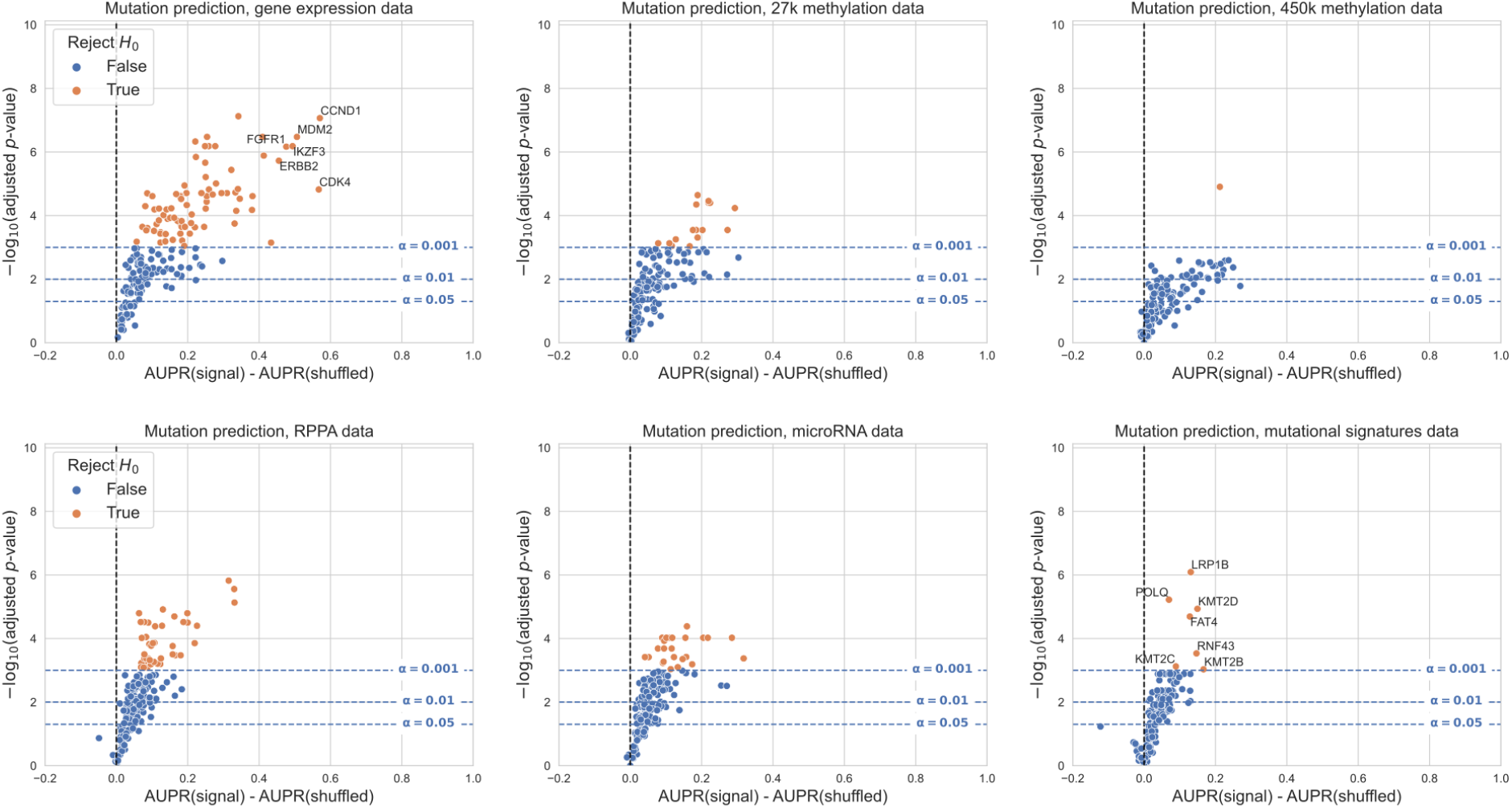
Volcano-like plots showing predictive performance for each gene in the cancer-related gene set for all data types, relative to the permuted baseline model, when genes are filtered based on the entire dataset rather than by cancer type. For this filtering approach, we included/excluded entire genes rather than individual cancer types: specifically, we trained a classifier for each gene where all cancer types combined had at least 5% mutated samples and at least 100 total mutated samples, resulting in 182 total classifiers. The *x*-axis shows the difference in mean AUPR compared with a baseline model trained on permuted labels, and the *y*-axis shows *p*-values for a paired *t*-test comparing cross-validated AUPR values within folds. Counts of genes making the significance threshold of 0.001: gene expression 81/182 (44.5%), 27K methylation 16/182 (8.8%), 450K methylation 1/182 (0.6%), RPPA 41/182 (22.5%), microRNA 25/182 (13.7%), mutational signatures 7/182 (3.9%).

**Figure 12:**
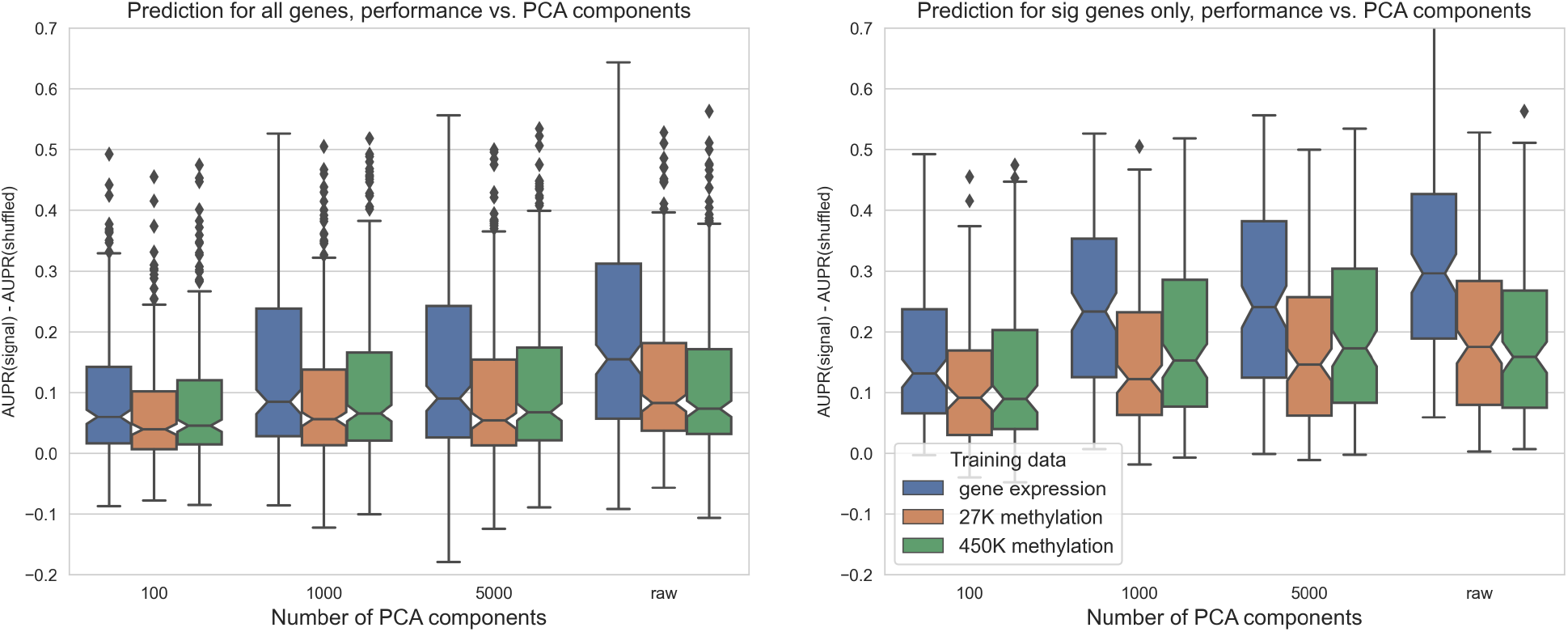
Predictive performance for genes in the cancer-related gene set, using each of the three data types as predictors. The *x*-axis shows the number of PCA components used as features, “raw” = no PCA compression.

**Figure 13:**
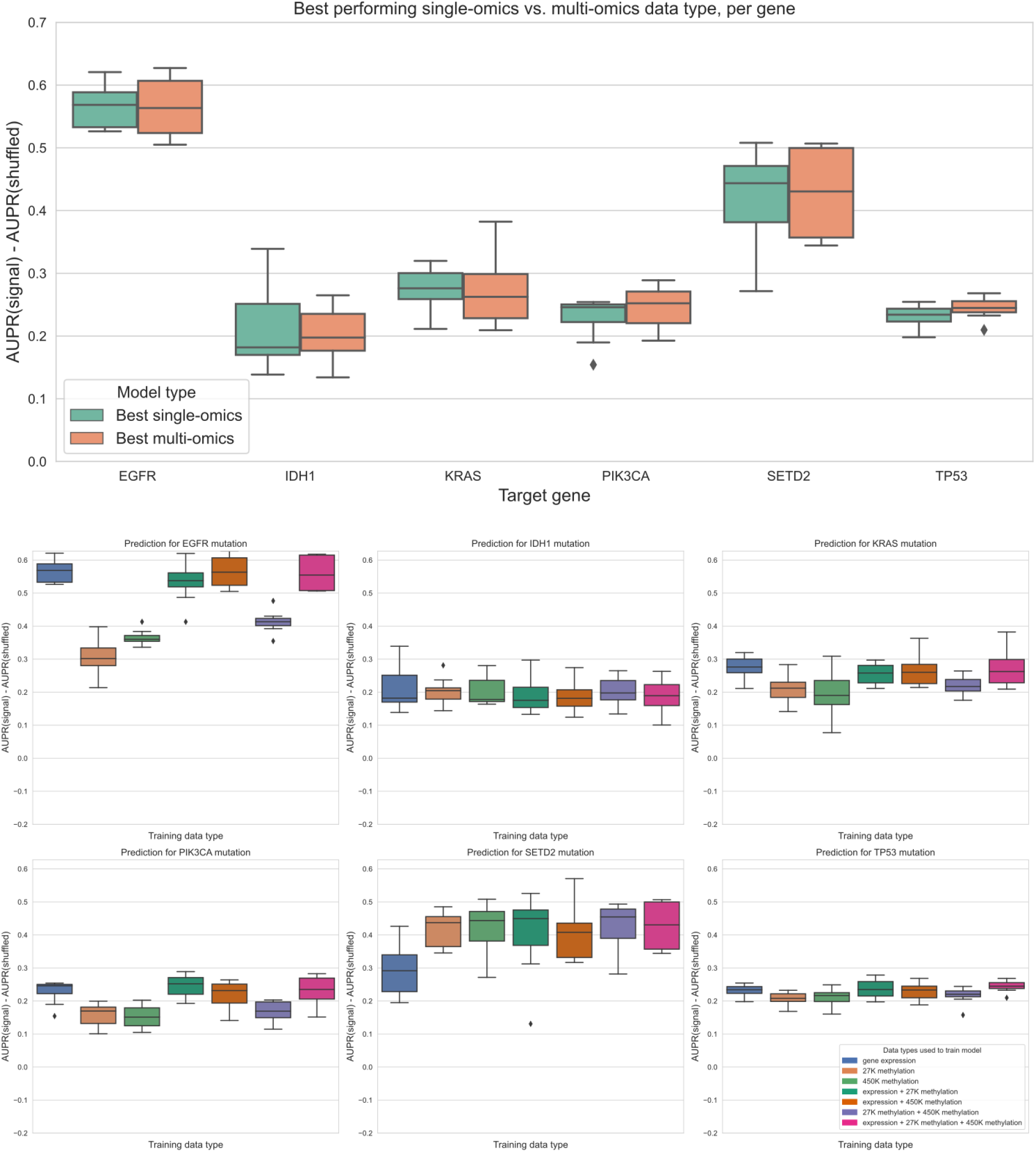
Top plot: comparing the best-performing model (i.e. highest mean AUPR relative to permuted baseline) trained on a single data type against the best “multi-omics” model for each target gene, using raw (not PCA compressed) features. For feature parity between data types, the top 20,000 features by mean absolute deviation were used for each data type. The difference between single-omics and multi-omics performance for *TP53* was statistically significant (*p*=0.0078), but other differences between single-omics and multi-omics models were not statistically significant using paired-sample Wilcoxon tests across cross-validation folds, for a threshold of 0.05. Bottom plots: classifier performance, relative to baseline with permuted labels, for individual genes. Each panel shows performance for one of the six target genes; box plots show performance distribution over 8 evaluation sets (4 cross-validation folds x 2 replicates).

**Figure 14:**
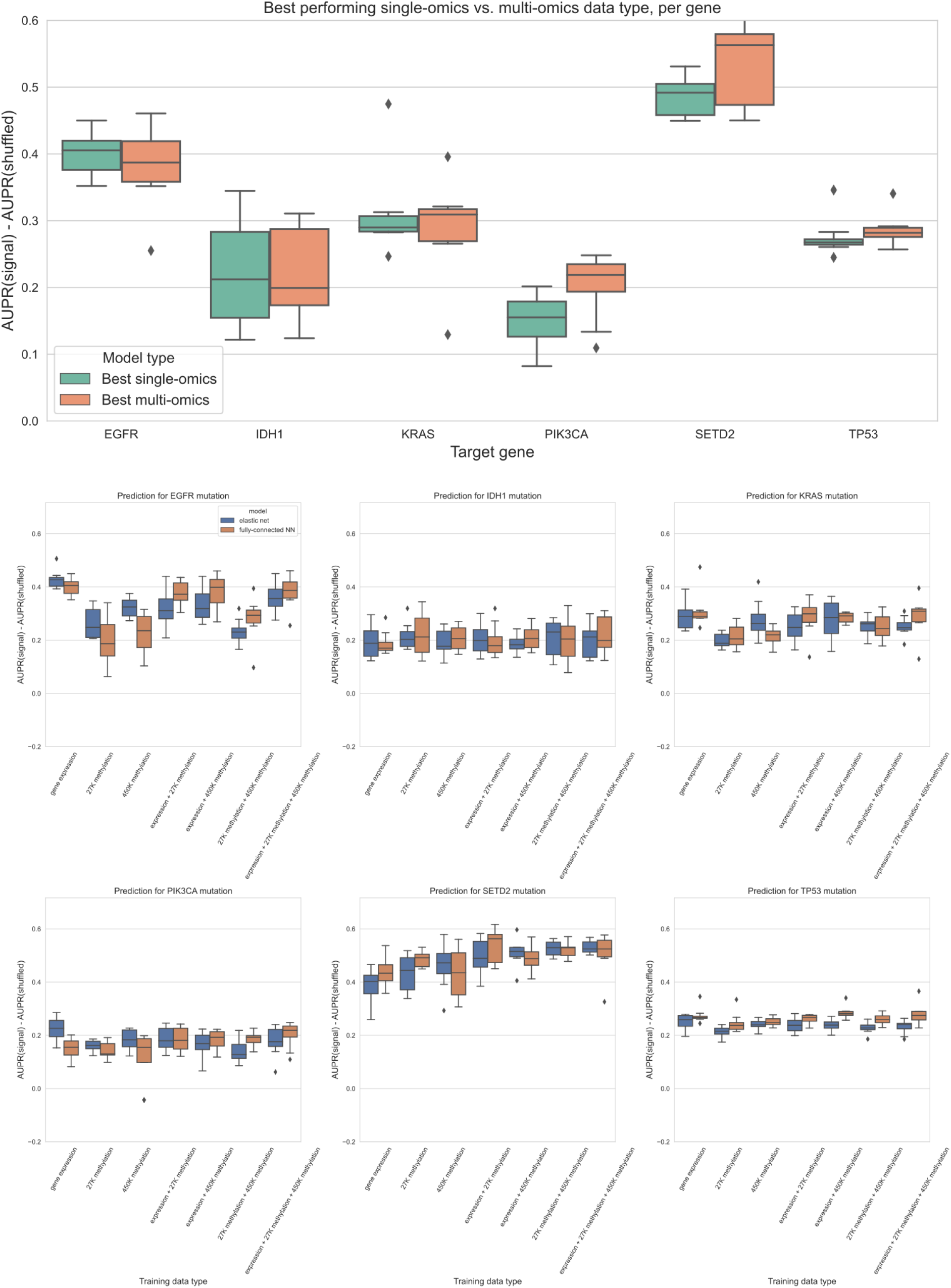
Top plot: comparing the best-performing model (i.e. highest mean AUPR relative to permuted baseline) trained on a single data type against the best “multi-omics” model for each target gene, using a 3-layer fully-connected neural network. The top 5,000 principal components were used as predictive features for each data type. The difference between single-omics and multi-omics performance for *PIK3CA* (*p* = 0.0156, in favor of multi-omics) and *TP53* (*p* = 0.0391, in favor of single-omics) were statistically significant, but other differences between single-omics and multi-omics models were not statistically significant using paired-sample Wilcoxon tests across cross-validation folds, for a threshold of 0.05. Bottom plots: comparison of classifier performance between elastic net and fully-connected NN, relative to baseline with permuted labels, for individual genes. Each panel shows performance for one of the six target genes; box plots show performance distribution over 8 evaluation sets (4 cross-validation folds x 2 replicates).

## Discussion

We carried out a large-scale comparison of data types in the TCGA Pan-Cancer Atlas as functional readouts of genetic alterations in cancer, integrating results across cancer types and across driver genes. Overall, we found that gene expression captures signatures of mutation state most effectively in general, relative to a baseline model, but we saw that for many genes other data types could be equally effective at predicting mutation presence or absence. For pan-cancer survival prediction, we found that the functional readouts tended to be similarly effective, outperforming a simple baseline using age and sample mutation burden in most cases. Our multi-omics modeling experiment indicated that the mutation state information captured by gene expression and DNA methylation is highly redundant, as added data types resulted in no gain or modest gains in classifier performance.

Comparing mutation status prediction using raw and PCA compressed expression and DNA methylation data, we observed that feature extraction using PCA provided no benefit compared to using raw gene or CpG probe features. Other studies using DNA methylation array data have found that nonlinear dimension reduction methods, such as variational autoencoders and capsule networks, can be effective for extracting predictive features [58,59]. The latter approach is especially interesting because capsule networks and “capsule-like methods” can be constrained to extract features that align with known biology (i.e. that correspond to known disease pathways or CpG site annotations).

This can improve model interpretability as well as predictive performance. Similar methods have been applied to extract biologically informed features from gene expression data (see, for instance, [60,61]). A more comprehensive study of dimension reduction methods in the context of mutation prediction, including the features selected by these methods and their biological relevance and interpretation, would be a beneficial area of future work. In addition to methods for extracting features, another aspect of the study that could be explored further is methods for labeling samples as mutated or not more eiciently. Although the mutation calls we used from MC3 represent the consensus of multiple algorithms aggregated through a standard pipeline, benchmarking other methods for identifying mutated samples could improve the utility of our method, such as calling mutations directly from RNA-seq data to avoid the need for paired samples [62,63].

In contrast to many other studies demonstrating the benefits of integrating multiple -omics data types for various cancer-related prediction problems [64,65,66,67,68], we found that combining multiple data types to predict mutation status was generally not effective for this problem. The method we used to integrate different data types by concatenating feature sets is sometimes referred to as “early” data integration (discussed in more detail in [69] and [70]). It is possible that more sophisticated data integration methods, such as “intermediate” integration methods that learn a set of features jointly across datasets, would produce improved predictions. We do not interpret our results as concrete evidence that multi-omics integration is not effective for this problem; rather, we see them as an indication that this is a challenging data integration problem for which further investigation is needed. We also present this problem as a set of benchmark tasks on which multi-omics integration methods can be evaluated. In addition to the methodological questions, the issue of data integration also has implications for the underlying biology: a more nuanced understanding of when different data readouts provide redundant information, and when they can contribute unique information about cancer pathology and development, could have many translational applications.

One limitation of the current study is that, for the mutation prediction problem, we only evaluated classifiers that were trained on pan-cancer data. Considering every possible combination of target gene and TCGA cancer type (85 target genes x 33 cancer types x 6 data types) would have drastically increased the computational load and presented a large multiple testing burden. Alternatively, choosing only a subset of gene/cancer type combinations to study would have biased our results toward known driver gene/cancer type relationships, which we aimed to avoid. In future work it would be interesting to identify classifiers that perform well in a certain cancer type but not in the pan-cancer context and to compare these instances across different cancer types. As a motivating example, other studies have shown that activating mutations in Ras isoforms (*HRAS, KRAS, NRAS*) tend to have similar effects to one another in thyroid cancer, producing similar gene expression signatures [15]. In multiple myeloma, however, activating *KRAS* and *NRAS* mutations produce distinct expression signatures, necessitating separate classifiers [71]. A high-throughput computational pipeline to identify cases where functional signatures of a particular cancer driver are either concordant or discordant between cancer types could identify opportunities for context-specific protein function prediction, improve biomarker identification, and suggest cases where drugs targeting specific alterations might produce discordant results in different cancer types.

As with any study relying on observational, cross-sectional data such as the TCGA Pan-Cancer Atlas, the conclusions that we can draw are limited by the data. In particular, for any of our “well-predicted” genes (i.e. genes that, when mutated, have strong signatures in one or more data types), we cannot definitively distinguish correlation from causation. To directly assess the effects of particular mutations on various data modalities, some studies use cell line data from sources such as the Cancer Cell Line Encyclopedia (CCLE) [72]. While this approach could help to isolate the causal effect of a given mutation on a given cell line, cell lines are sometimes an imperfect match for the cancers they are derived from [73]. We are also limited in that we cannot assign timing or clonal status to mutations, or fully characterize intratumor heterogeneity, with certainty from the bulk sequencing data generated by TCGA (although some features of tumor mutational processes over time can be estimated from bulk data, e.g. [74]). As methods for generating large longitudinal datasets at single-cell resolution mature and scale, we will need to revise the way we think about cellular function and dysregulation in cancer cells, as dynamic and adaptive processes rather than a single representative snapshot of a tumor.

Based on our results, for studies focused on the functional consequences of cancer mutations, we recommend that researchers cancers prioritize downstream readouts based on the gene or genes of interest (Figure 6). On balance, prediction of mutation status is best in general using gene expression data, and prediction of patient survival is similar for all data types in the study. However, the finding that for many genes, multiple functional profiles contain much of the same information will be useful for some study designs, given varying cost and stability of different readouts. In addition to gene expression, results using DNA methylation and RPPA measurements as predictive features were promising, especially considering the substantially lower dimensionality of the RPPA dataset compared to other data types. It is important to note that the specific technologies chosen by TCGA, and the tradeoffs inherent in such a high-throughput study, could influence the comparison: it is possible that, for instance, another technology for measuring DNA methylation (such as bisulfite sequencing) or another technique for measuring protein abundance (such as mass spectrometry-based proteomics) could change performance for those data types. Future technology advances, in both quality and quantity of data, are likely to improve our understanding of the full picture of functional consequences of mutations in cancer cells.

## Declarations

### Ethics approval and consent to participate

Not applicable

### Consent for publication

Not applicable

### Availability of data and materials

The datasets analyzed during this study were previously published as part of the TCGA Pan-Cancer Atlas project, and are publicly available from the NIH NCI Genomic Data Commons (GDC) at https://gdc.cancer.gov/about-data/publications/pancanatlas.

Scripts used to download and preprocess the datasets for this study are available at https://github.com/greenelab/mpmp/tree/master/00_download_data.

## Authors’ contributions

JC: conceptualization, methodology, software, visualization, writing - original draft, writing - review and editing BCC: methodology, writing - review and editing MC: methodology, writing - review and editing CSG: conceptualization, funding acquisition, methodology, supervision, writing - review and editing

### Acknowledgements

We would like to thank Alexandra Lee, Ariel Hippen, Ben Heil, Milton Pividori, and Natalie Davidson for reviewing the software associated with this work and providing insightful feedback. Figure 1 (the schematic of the background and evaluation pipeline) was created using BioRender.com.

## Supplementary Information

A version of the main paper figures using the area under the receiver-operator curve (AUROC) metric rather than AUPR is available at https://doi.org/10.6084/m9.figshare.14919729.

In a previous version of this paper, we ran our analysis only for the genes in the Vogelstein et al. [38] gene set. While there were some gene-to-gene differences in this set, we did not observe large differences between methylation and gene expression performances overall. Scaling up the gene set by combining cancer gene sets from the literature as described in the methods/results sections affected the study results somewhat, as mutations in the added genes tend to be better predicted using gene expression than other data types. During the revision, we explored the difference between the genes in this gene set and the genes in the “merged” cancer-related gene set but not in the Vogelstein genes. GO analysis results for the Vogelstein genes are available at https://doi.org/10.6084/m9.figshare.19565890, and results for the non-Vogelstein genes are available at https://doi.org/10.6084/m9.figshare.19565887. We noticed that the non-Vogelstein genes tend to be enriched for terms relating to transcription factors and transcriptional regulation.

As a data resource, coeicients and hyperparameter choices for final models fit using gene expression features are available on Figshare: coeicients are available at https://doi.org/10.6084/m9.figshare.19576012 and hyperparameters are at https://doi.org/10.6084/m9.figshare.19576048. Columns in the coeicients dataset correspond to target genes (gene symbols), and rows correspond either to gene expression features (Entrez IDs) or covariates (cancer type indicator variables or log(mutation burden)). An ‘NA’ value in a cell indicates that feature was not used in the model for the corresponding gene (for a gene feature this means that gene was not in the top 8000 genes by MAD, for a cancer type feature this means that cancer type was not included in the training set based on our mutation filters). A 0 value in a cell indicates that feature was included in model training, but it was not selected by the elastic net feature selection algorithm. Columns in the hyperparameters dataset correspond to hyperparameters (alpha and l1_ratio for elastic net logistic regression) and rows correspond to target genes.

Regarding the hyperparameters for the final models, recall that for the main figures in the paper, we evaluate each of our models using 2 replicates of 4-fold cross-validation. For each of these folds (train/test splits), we further split the training set into train and validation sets to select hyperparameters, independently for each fold, and evaluate the models on the test set to get the results in the paper. Because we are evaluating performance over multiple folds, it is not perfectly straightforward to get a single set of regression coeicients, since we have a (potentially different) set of coeicients for each cross-validation fold. In order to synthesize these results into a single model for each gene in each data type, we selected one of the 8 sets of hyperparameters (from the 8 best models, 1 per CV fold) at random, with probability proportional to performance (AUPR) on the validation set used to select the hyperparameters, described above (so test set performance is not used here). We then used the selected hyperparameters to train a single model on the entire dataset.

**Table 1:**
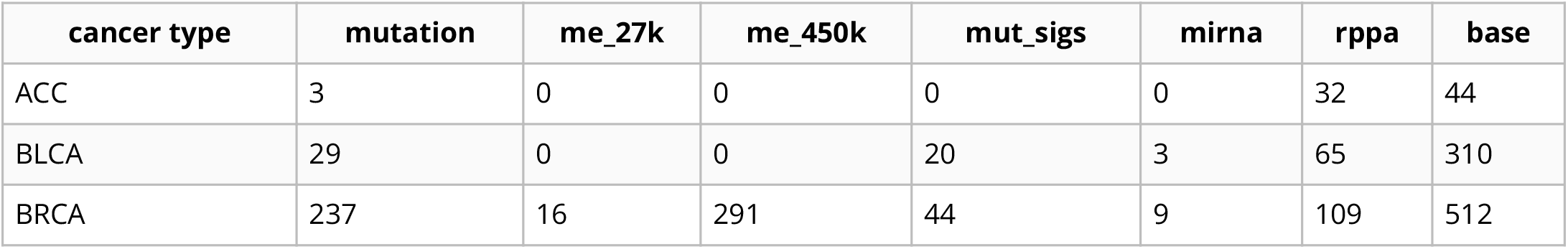
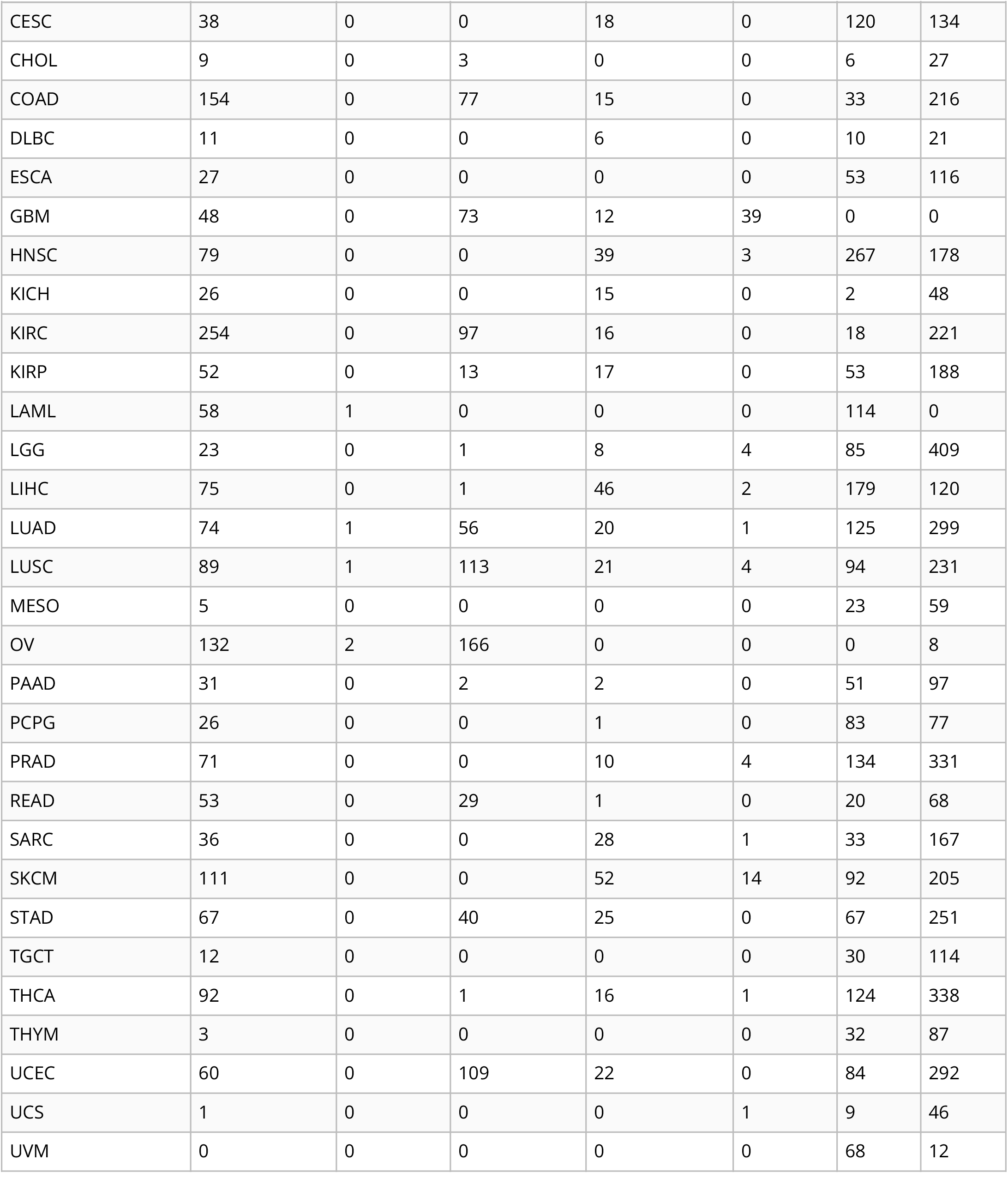
Number of samples from each TCGA cancer type that are “dropped” as more data types are added to the analysis. The “base” column indicates the number of samples that are present per cancer type in the final intersection of all data types (i.e. each sample counted in the last column has data for each of the 7 data types, including gene expression (not listed here) and somatic mutations).

